# Phenotypic deconvolution in heterogeneous cancer cell populations using drug screening data

**DOI:** 10.1101/2022.01.17.476604

**Authors:** Alvaro Köhn-Luque, Even Moa Myklebust, Dagim Shiferaw Tadele, Mariaserena Giliberto, Leonard Schmiester, Jasmine Noory, Elise Harivel, Polina Arsenteva, Shannon M. Mumenthaler, Fredrik Schjesvold, Kjetil Taskén, Jorrit M. Enserink, Kevin Leder, Arnoldo Frigessi, Jasmine Foo

**Affiliations:** Oslo Centre for Biostatistics and Epidemiology, Faculty of Medicine, University of Oslo, 0372 Oslo, Norway; Department of Molecular Cell Biology, Institute for Cancer Research, Oslo University Hospital, 0379 Oslo, Norway; Centre for Cancer Cell Reprogramming, Institute of Clinical Medicine, Faculty of Medicine, University of Oslo, 0318 Oslo, Norway; Translational Hematology and Oncology Research, Cleveland Clinic, Cleveland, OH 44131, USA; Dept. of Cancer Immunology, Institute for Cancer Research, Oslo University Hospital, 0310 Oslo, Norway; KG Jebsen Center for B-Cell Malignancies, Institute for Clinical Medicine, University of Oslo, 0450 Oslo, Norway; Institute for Mathematics and its Applications, School of Mathematics, University of Minnesota, Minneapolis, MN 55455, USA; ENSTA, Institut Polytechnique de Paris, 91120 Palaiseau, Paris, France; Institut de Matématiques de Bourgogne, Universite de Bourgogne, 21078 Dijon Cedex, Dijon, France; Lawrence J. Ellison Institute for Transformative Medicine, Los Angeles, CA 90064, USA; Department of Biomedical Engineering, Viterbi School of Engineering, University of Southern California, Los Angeles, CA 90089, USA; Department of Oncology, Norris Comprehensive Cancer Center, Keck School of Medicine, University of Southern California, CA 90033, USA; Oslo Myeloma Center, Department of Hematology, Oslo University Hospital, 0450 Oslo, Norway; Section for Biochemistry and Molecular Biology, Faculty of Mathematics and Natural Sciences, University of Oslo, 0037 Oslo, Norway; College of Science and Engineering, University of Minnesota, Minneapolis, MN 55455, USA; Oslo Centre for Biostatistics and Epidemiology, Oslo University Hospital, 0372 Oslo, Norway

**Author notes:** These authors contributed equally to this work (Alvaro Köhn-Luque, Even Moa Myklebust). Department of Medical Genetics, Oslo University Hospital, Oslo, Norway (Dagim Shiferaw Tadele). These authors contributed equally to this work (Kevin Leder, Arnoldo Frigessi, Jasmine Foo).

## Abstract

Tumor heterogeneity is an important driver of treatment failure in cancer since therapies often select for drug-tolerant or drug-resistant cellular subpopulations that drive tumor growth and recurrence. Profiling the drug-response heterogeneity of tumor samples using traditional genomic deconvolution methods has yielded limited results, due in part to the imperfect mapping between genomic variation and functional characteristics. Here, we leverage mechanistic population modeling to develop a statistical framework for profiling phenotypic heterogeneity from standard drug screen data on bulk tumor samples. This method, called PhenoPop, reliably identifies tumor subpopulations exhibiting differential drug responses, and estimates their drug-sensitivities and frequencies within the bulk. We apply PhenoPop to synthetically-generated cell populations, mixed cell-line experiments, and multiple myeloma patient samples, and demonstrate how it can provide individualized predictions of tumor growth under candidate therapies. This methodology can also be applied to deconvolution problems in a variety of biological settings beyond cancer drug response.

**Motivation:** Tumors are typically comprised of heterogeneous cell populations exhibiting diverse phenotypes. This heterogeneity, which is correlated with tumor aggressiveness and treatment failure, confounds current drug screening efforts aimed at informing therapy selection. In order to optimally select treatments, understanding the frequency and drug-response profile of individual subpopulations within a tumor is critical. Furthermore, quantitative profiles of tumor drug-response heterogeneity, in combination with predictive mathematical modeling of tumor dynamics, can be used to design effective temporal drug-sequencing strategies for tumor reduction.

Here, we present a method that enables the deconvolution of tumor samples into individual sub-components exhibiting differential drug-response. This method relies on standard bulk drug-screen measurements and outputs the frequencies and drug-sensitivities of tumor subpopulations. This framework can also be applied more broadly to deconvolve cellular populations with heterogeneous responses to a variety of external stimuli and environmental factors.

## Introduction

Most human tumors display a striking amount of phenotypic heterogeneity in features such as gene expression, morphology, metabolism, and drug response. This diversity fuels tumor evolution and adaptation, and it has been correlated with higher risks of treatment failure and tumor progression.^1–8^ Indeed, treatments that initially elicit clinical response can select for drug-tolerant tumor subpopulations, leading to outgrowth of resistant clones and tumor recurrence. Additionally, the heterogeneity and composition of tumors is known to vary widely between patients, underscoring the need for more personalized approaches to cancer therapy that profile and address intra-tumor heterogeneity and its evolutionary consequences. Towards this goal, recent advances in single-cell genomic profiling of tumor samples have enabled the assessment of the genetic variability within tumor cell populations. However, single-cell technologies are often limited by large measurement errors, incomplete coverage, and small sample availability, which leads to challenges in capturing the temporal dynamics crucial for understanding response to therapies. Furthermore, the mapping between genotypic and phenotypic variation is far from perfect: not all variation in cellular drug response can be explained by genetic mechanisms, and divergent genetic profiles can lead to similar treatment responses.^9,10^

Another important approach to designing individualized treatment strategies is personalized drug sensitivity screening, a procedure in which patient tumor samples are tested for functional responsiveness to a library of drugs using high throughput *in vitro* drug sensitivity assays. In these assays, cells are treated with various concentrations of a drug and the number of viable cells is measured at one or more fixed time points. The resulting data are normalized and fitted to produce viability curves, whose summary characteristics (e.g. *IC*_50_, *EC*_50_,*AUC*) are used to compare drug sensitivity across multiple drugs and/or cell populations.^11–15^ Increasingly, such drug screens are used as a tool in personalized medicine to evaluate and rank the potential efficacy of therapeutic agents on a patient’s disease cell population. However, the interpretation of these cell viability curves and associated metrics are confounded by the presence of cellular heterogeneity within the population. In particular, the presence of multiple subpopulations with divergent drug response characteristics may result in an intermediate drug sensitivity profile that does not accurately represent any individual cell type within the population.^16^ Developing techniques to detect the presence of subpopulations with distinct drug sensitivity profiles is crucial for achieving effective treatment strategies.

In this work, we develop a methodology for detecting the presence of cellular subpopulations with differential drug responses, using standard bulk cell viability assessment data from drug screens. Our method, PhenoPop, detects the presence and composition fractions of distinct phenotypic components in the tumor sample and quantifies the sensitivity of each subpopulation to a specific mono-therapy. Notably, since the cell counts used in the deconvolution are aggregated signals from the entire population, the approach is categorically different from previous techniques that rely on observations at the individual level such as growth mixture modeling^17^ or latent class growth analysis.^18^ PhenoPop utilizes statistical tools in combination with an underlying population dynamic model describing the evolution of a heterogeneous mixture of tumor cells with differential drug sensitivity over time. In this work, we validate PhenoPop using simulated tumor drug screening data as well as measurements of drug response in known mixture experiments of cancer cell lines. We then use PhenoPop to profile the population drug response heterogeneity in multiple myeloma patient samples, and we demonstrate how these results can be used to produce personalized predictions of tumor response to therapy. This methodology can be applied across a variety of cancer types and therapies to characterize the drug-response heterogeneity within tumors.

## Results

Figure 1 provides an overview of the PhenoPop workflow. First, a tumor sample is extracted, divided, and exposed to a panel of therapeutic compounds at a range of concentrations. For each drug, the population size counts are measured at a series of time points for each concentration and replicate. This data is then used as the input to PhenoPop, which estimates the parameters of the underlying population dynamic model for each candidate number of subpopulations. Then, a model selection process is performed to identify the number of subpopulations present and to estimate the mixture fractions and drug sensitivities of each subpopulation. Details are provided in the STAR methods section.

**Figure 1:**
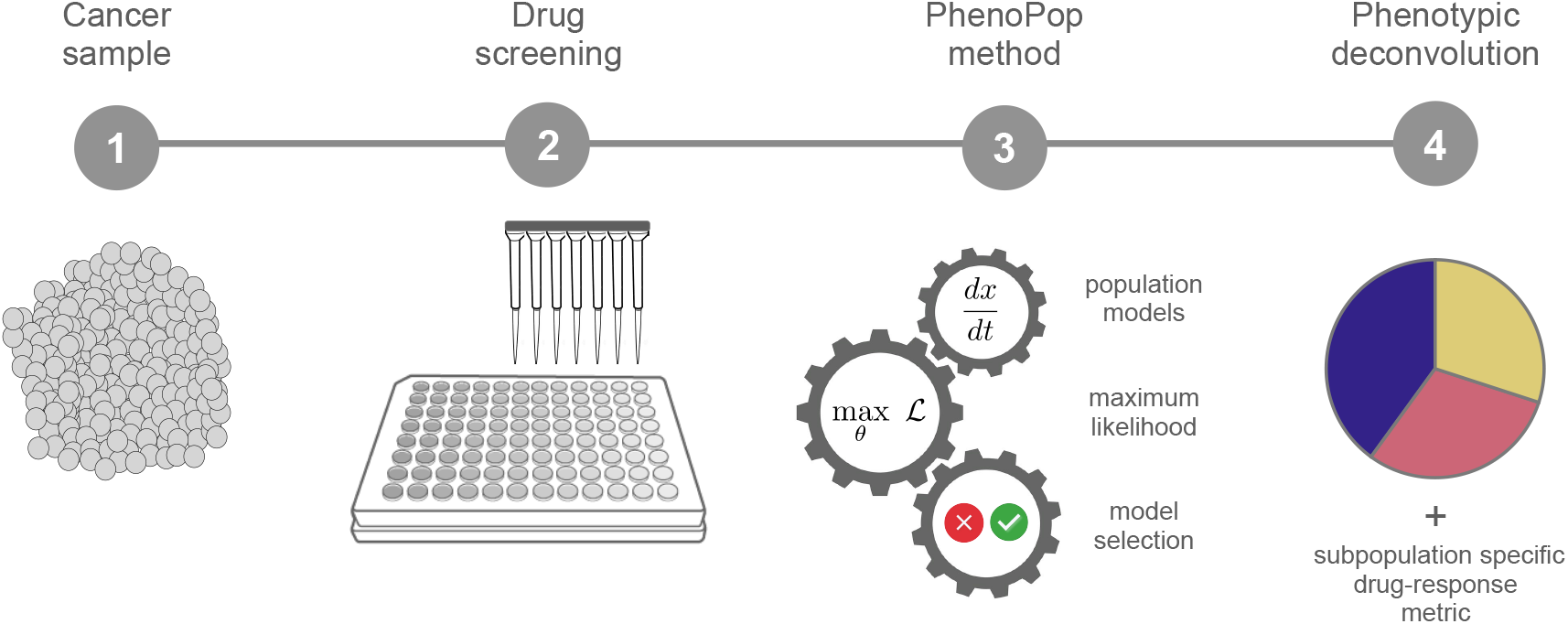
The PhenoPop workflow: 1) A cancer sample is taken from a patient. 2) Drug screening is performed on the bulk sample. 3) Population deconvolution is performed using PhenoPop. 4) Resulting identification of population subcomponents, their mixture fractions and drug-sensitivity.

### Validation in synthetic populations

To quantify the performance of PhenoPop in mixtures of 1, 2, and 3 populations, three synthetic populations were designed to have drug-response properties similar to cell lines observed in in-vitro experiments. Using the model for data generation described in the STAR methods section (subsection *Generation of synthetic population data*), synthetic data was generated for 9 different mixture compositions of the three populations. The synthetic mixtures were exposed to 17 concentrations of the simulated drug, and the bulk cell populations were measured at 9 equidistant points in time. The simulated drug concentrations were chosen to cover the range where population growth rates were affected by changes in the drug concentration. To simulate measurement error, random noise was added to each bulk cell count. Data from 4 replicates of the experiment were used to perform the inference.

To measure drug sensitivity, PhenoPop uses the growth rate-associated metric *GR_50_*, introduced in Hafner et al.^19^ and defined as the concentration at which the population growth rate is reduced by half of the maximum observed effect, as it provides a robust metric for comparing drug-response across cellular subpopulations (STAR methods, subsection *Calculation of GR*_50_ *values*). To assess the accuracy of the PhenoPop deconvolution analysis, (i) the estimated mixture fractions from the deconvolution were compared with the true mixture fractions, and (ii) the GR50 obtained from the deconvolution was compared with the true GR50 region. The use of a region, or range of values, for the true GR50 reflects the inherent limitation from sampling discrete concentrations in experimental data; it is only possible to ascertain that the GR50 is somewhere between the closest two sampled concentration levels, and the finer the sampling resolution, the smaller the range of uncertainty.

Figure 2A shows true mixture compositions and GR50 values compared to PhenoPop’s estimates for the 9 cases, in an experiment where the noise terms were sampled independently from a Gaussian distribution with mean 0 and base noise level of 5 %, meaning that the standard deviation of the noise terms equaled 5 % of the noiseless cell count at time 0. Additional sensitivity tests evaluating PhenoPop performance on synthetic data with varying noise levels (up to 50 %) are discussed in the section *PhenoPop experimental design recommendations* and data are provided in the Supplemental Information. To place these noise levels in the context of expected noise levels from experimental drug screen data, the standard deviation to mean ratio reported from several common automated or semi-automated cell counting techniques ranges from 1-15 %.^20,21^ For example, counts obtained via a trypan blue exclusion-based Vi-CELL® XR Cell Viability Analyzer (Beckman Coulter) had noise levels consistently less than 5.3 % across several cell lines,^20^ while those obtained via a Countess® Automated Cell Counter (Invitrogen) fell in the range 11-14.3 %. Cells counts obtained using the Cellomics ArrayScan high content screening platform in another set of experiments (used in this work) had standard deviation to mean ratios of 1-5.6 %.^21^

**Figure 2:**
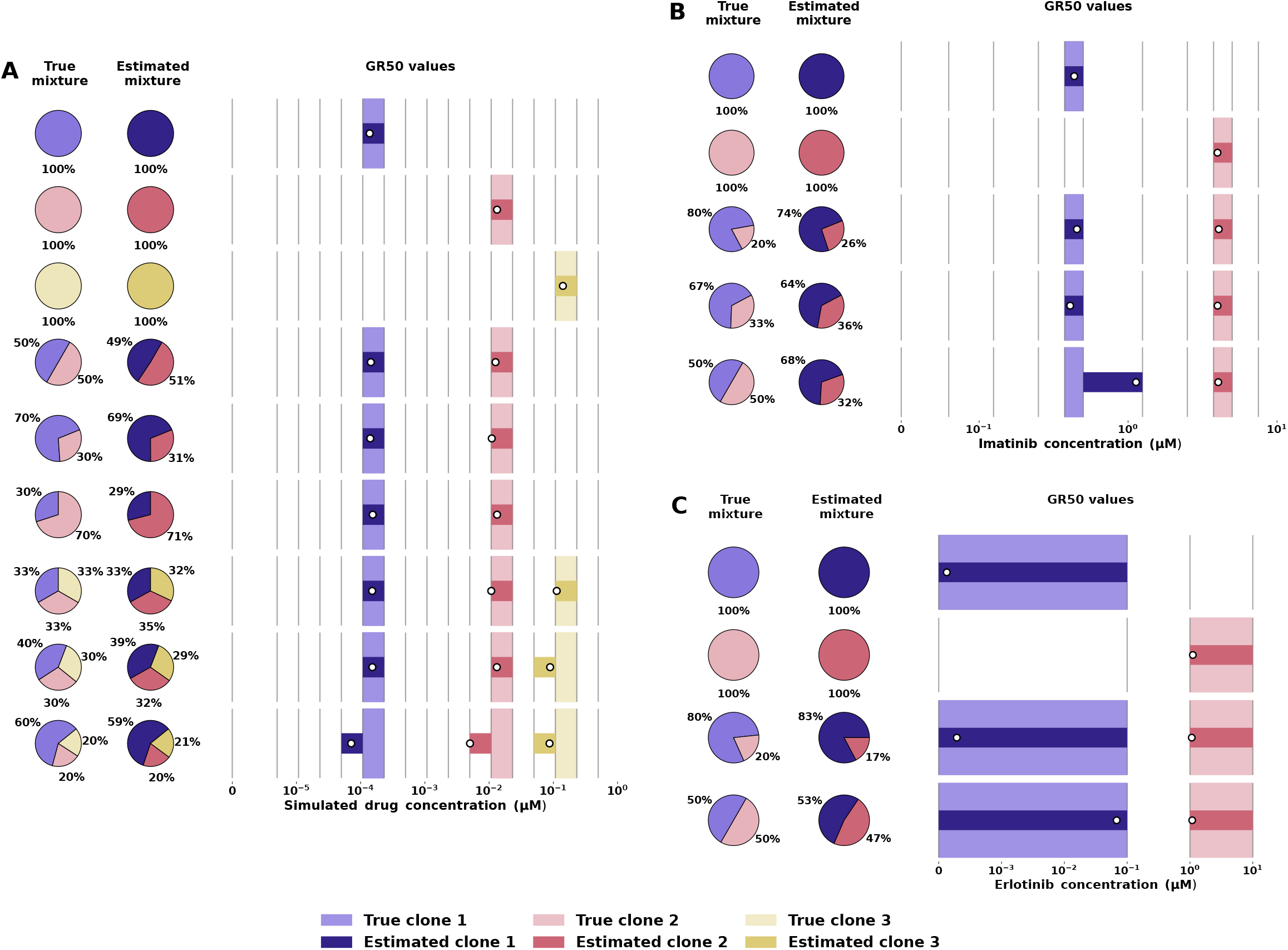
Validation of the PhenoPop method. Deconvolution results in A: simulated data, B: Ba/F3 murine cell-line derived experimental data and C: NSCLC cell-line derived experimental data. In each of A, B, C, each row corresponds to an independent experiment. On the left, the true and estimated mixture percentages are shown. On the right, the grid of vertical lines corresponds to drug concentrations. The true GR50 region is marked by a light-colored bar extending between two adjacent concentrations, and the estimated GR50 region is marked by a narrower darker-colored bar. The point estimate of the GR50 is marked with a white circle. Details of model selection results are shown in Supplemental Figure S3.

Figure 2A demonstrates that PhenoPop inferred the mixture fractions within 2 percentage points for mixtures of 1, 2, and 3 populations at the 5 % noise level. The GR50 values were inferred precisely within the true GR50 region for all mixtures of 1 and 2 populations, and also for an equal mixture of 3 populations. In the case with 3 populations in a 40:30:30 mixture, one of the estimated GR50 values is off by 1 GR50 region, and in the 3-population mixture with a 60:20:20 mixture, all three estimated GR50 values are off by 1 GR50 region. Details of model selection results are shown in Supplemental Figure S3A.

### Validation with cell line experiments

Next, to investigate the performance of our method in the experimental setting, mixtures of cell populations with differential drug sensitivity were constructed and subjected to drug screen experiments. The resulting bulk cell population readings at varying drug concentrations, time points, and replicates were used as inputs to PhenoPop.

#### Imatinib-sensitive and -resistant Ba/F3 cells

We tested monoclonal and mixture populations of isogenic Ba/F3 murine cell lines that were stably transformed with either the wild-type BCR-ABL fusion oncogene or with BCR-ABL-T315I, which contains a point mutation that confers increased resistance to the Abl tyrosine kinase inhibitor imatinib. Note that expression of these oncogenes renders cells addicted to BCR-ABL activity.^22^ Monopopulations and mixtures of these two cell lines were treated with 11 different concentrations of imatinib, and the bulk cell population sizes were quantified at 14 time points. Using this bulk population data, PhenoPop was able to correctly assess the number of component subpopulations (Figure 2B). As shown in Figure 2B, PhenoPop also estimated the fraction of the population belonging to each subcomponent at the start of the drug screen as well as the drug sensitivity (GR50) of each subpopulation; the estimates demonstrated good agreement with the known mixture proportions and independently assessed GR50 ranges of the monoclonal T315I+/- populations. Details of model selection results are shown in Supplemental Figure S3B.

#### Erlotinib-sensitive and -resistant NSCLC cells

Additionally, two EGFR-mutant non-small cell lung cancer (NSCLC) lines, HCC827 and H1975, were considered for their differential sensitivity to the drug compound erlotinib. The mutation T790M, which is present in H1975 cells but not in HCC827 cells, confers increased resistance to erlotinib. Monopopulations and mixtures of the erlotinib-sensitive and -resistant NSCLC cell lines were treated with four drug concentrations and total cell population count was assessed at 0, 24, 48, and 72 hours with four replicates.^21^ Figure 2C demonstrates PhenoPop’s results on this bulk data. PhenoPop was able to correctly assess when populations were monoclonal, as well as to detect the presence of two populations in the bulk drug response data from mixed populations. Furthermore, using the bulk mixture response data, PhenoPop accurately estimated the mixture fractions and GR50 values of each component subpopulation. The reference GR50 ranges were independently assessed on monoclonal HCC827 and H1975 cell populations. Details of model selection results are shown in Supplemental Figure S3C.

### Deconvolution analysis of Multiple Myeloma patient samples

Next, PhenoPop was used in a clinical scenario to deconvolve twenty drug sensitivity screens performed on five Multiple Myeloma (MM) patient samples. MM is a clonal B-cell malignancy characterized by abnormal proliferation of plasma cells in the bone marrow. The median survival time of MM patients is about 6 years, with a disease course typically marked by multiple recurrent episodes of remission and relapse.^23^ Drug responses and relapses are currently unpredictable, largely due to unknown complex clonal compositions and dynamics under treatment.^24,25^

Bone marrow samples were taken from each patient, processed, and screened with a set of MM clinically-relevant drugs, as illustrated in Figure 3A and described in the STAR methods section.^26^ To perform the drug screens, samples from each patient were subjected to treatment at varying concentrations with a subset of the following drugs: Dexamethasone, Ixazomib, Melflufen, Selinexor, Thalidomide, and Venetoclax. We note that screening data for all drugs for each patient was not available; Figure 3B-G shows the set of patient samples treated by each drug and summarizes the results of PhenoPop deconvolution analysis on each set of drug screen data. The results provide each sample’s population heterogeneity profile with respect to each screened drug; these profiles are typically expected to be distinct between therapies since the characteristics driving response to specific drugs may vary.

**Figure 3:**
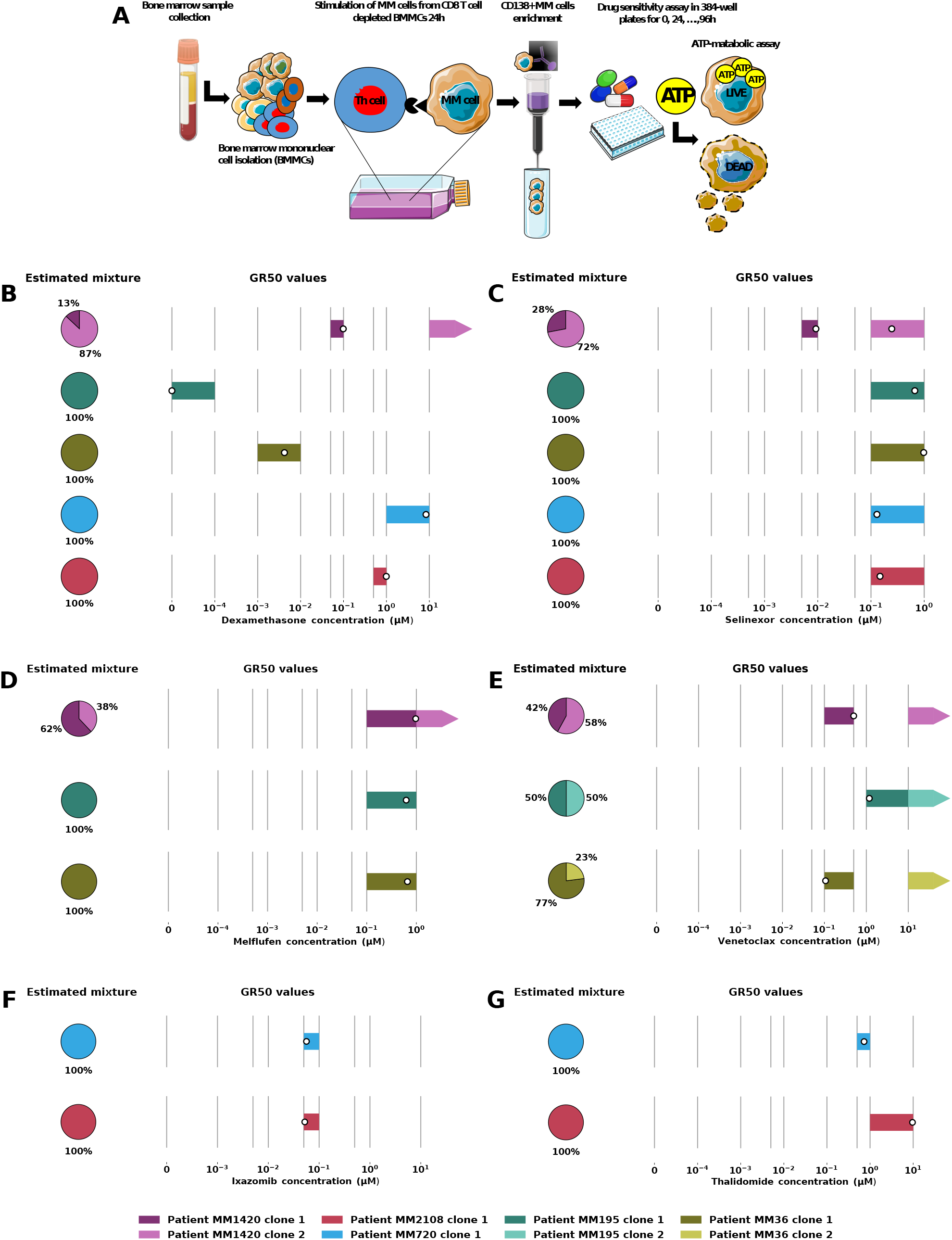
Phenotypic deconvolution of drug screens from MM patient samples. A: Illustration of the experimental protocol described in Methods^26^, created using smart.servier.com and biorender.com. B-G: Deconvolution results for 5 MM samples, each exposed to 4 out of 6 drugs: B: Dexamethasone, C: Selinexor, D: Melflufen, E: Venetoclax, F: Ixazomib, G: Thalidomide. In B-G separately, each row corresponds to a patient sample: On the left, the estimated mixture percentages are shown. On the right, the grid of vertical lines corresponds to drug concentrations. The estimated GR50 region is shown by a colored box, and the point estimate of the GR50 is marked with the white circle. If the inferred GR50 value of a population is above the highest observed concentration, the estimated GR50 is instead marked by an arrow pointing towards the right from the highest observed concentration.

#### Inter-patient similarities in subpopulation GR50s

In all cases PhenoPop identified either one or two subpopulations; details of model selection results are shown in Supplemental Figure S4. For example, Figure 3B shows that for patient MM1420, PhenoPop estimates that 87 % of the cells are resistant to Dexamethasone. This matches the clinically observed response, as the patient was refractory to Dexamethasone treatment in vivo. Interestingly, for all drugs used except Dexamethasone, the inferred subpopulations across patient samples share comparable GR50 values, although the proportions of these subpopulations may vary between patients (see Figure 3C-G). For example, for three patient samples treated with Venetoclax, PhenoPop inferred one more-sensitive and one more-resistant population (Figure 3E). However, the estimated proportions of the more-resistant populations (shown in the plot by the right-pointing arrows) varied from 23% up to 58%.

We hypothesized that subpopulations with similar GR50s across patients may in some cases be driven by similar genetic alterations. To investigate this, we also characterized the samples with inferred heterogeneous compositions for the presence of high-risk genomic abnormalities, including Gain(1q21) (2/3) and several mutations co-existing in the same screened sample (MM36). Interestingly, we noticed that the proportion of MM cells from two samples (MM1420 and MM195) harboring the aberration gain (1q21) (approximately 50 %) was similar to the PhenoPop-inferred mixture fractions for the more-resistant clone in the same two samples (50% and 58%, respectively). This supports our hypothesis that these subpopulations, which have similar levels of drug tolerance in different patients, may be driven the same alterations, and it is consistent with previous findings showing Gain(1q21) as negative predictor for Venetoclax efficacy in MM. This analysis provides genetic evidence that supports PhenoPop’s ability to profile phenotypic drug response heterogeneity.

#### Treatment response prediction using PhenoPop estimates

The utility of these phenotypic deconvolutions as initial states for predicting and optimizing patientspecific treatment schedules remains to be systematically explored. Here, as a proof of concept, we present a mathematical model to illustrate how to use the PhenoPop estimates of population frequencies and differential drug sensitivities to predict the treatment outcome for the three patients exposed to Venetoclax. For easier comparison, we assumed that all three patients start with a total of 10^12^ abnormal plasma cells. Figure 4 demonstrates how the same treatment dose, 2 μM of Venetoclax, assumed constant over the simulation for simplicity, leads to highly disparate treatment outcomes in patients with distinct phenotypic heterogeneity profiles uncovered by PhenoPop. In particular, we note that to observe the predicted relapse in patient MM36, simulations have to be run for a much longer time (3000 days) than for the other two patients. We also note that according to equation 3 the growth rate of a population can be negative at drug concentrations below the GR50. This is the case in the first panel of Figure 4, which shows both populations for patient MM1420 responding to Venetoclax at 2 μM while in Figure 3B the largest subpopulation is estimated to have a GR50 value that is above 10 μM. See the STAR methods section for a description of the model and its parameterisation.

**Figure 4:**
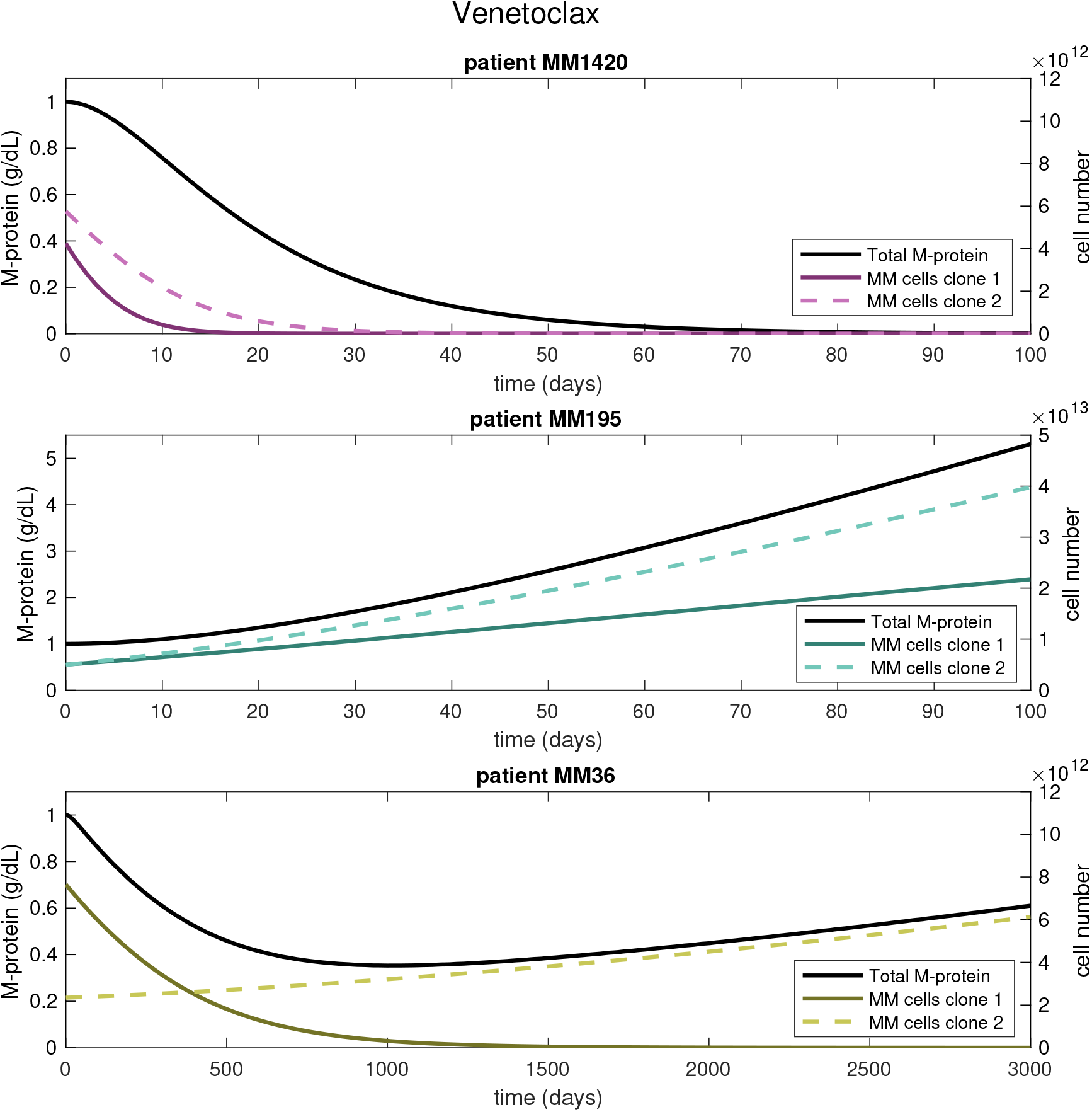
Proof-of-concept modeling of Multiple Myeloma disease dynamics under Vene-toclax treatment for three patients using PhenoPop deconvolution results. The estimated mixture and drug-response parameters obtained by PhenoPop (see Figure 3E) define the initial percentage of cells and drug-response for each clone and patient. Cells from both clones are assumed to produce monoclonal protein (M-protein), which can be used as a proxy for tumor burden. For easier comparison, we assume that all three patients start with a total of 10^12^ abnormal plasma cells (cell number shown in the right y-axes) and 1*g/dL* M-protein (shown in the left y-axes). All three patients are exposed to 2 μM of Venetoclax. See the STAR methods section for a description of the mathematical model.

To make treatment recommendations in a clinical setting, population deconvolution and subsequent predictive modeling of the heterogeneous cell population’s response to therapy should be performed for all candidate drugs. Depending on clinical goals (e.g. lowest tumor burden at particular time horizon, fastest tumor reduction, or longest time to resistance), the optimal treatment can be selected based on the predicted modeling outcomes under each therapy. Supplemental Figure S2 demonstrates a comparison of predicted response to Venetoclax versus three additional drugs (Dexamethasone, Selinexor and Meflufen) for which drug-screens were performed on the same patient samples. In this case, Dexamethasone may be the preferred option for patients MM195 and MM36, while for patient MM1420 all drugs produce a similar response.

## Discussion

Understanding the phenotypic heterogeneity of human tumors, especially in terms of drug response, is essential in treatment planning and prognosis prediction. The optimization of treatment regimens is a long-standing area of research in the mathematical oncology community;^27–30^ however, the initial state of the tumor, which strongly influences optimal treatment strategies, is typically unknown. PhenoPop enables the detection of tumor subpopulations, as well as estimation of their frequencies and drug sensitivities based on drug screen data. The resulting deconvolved tumor profile can be fed, as an initial state, into mathematical models of tumor dynamics to predict treatment response (see Figure 4) and identify optimal treatment regimens.

Though similar in objective, deconvolution methods are categorically different from clustering methods such as growth mixture modeling, since the former uses bulk observations from the combined population, while the latter relies on observations on the individual level. The deconvolution problem is in fact closer in nature to the problem considered in blind source separation in digital signal processing, in which one attempts to recover individual source components from a mixture of signals (see e.g. Moulines et al., Comon et al.^31,32^). However, a key assumption in this classic problem is the independence of the constituent components, a restriction that is not needed for PhenoPop. Interaction between individual populations, e.g. due to resource limitation or phenotypic switching, can be incorporated within the PhenoPop framework (see STAR Methods subsection Model Extension to Interacting Populations). The mathematical structure used in PhenoPop can also be applied to perform deconvolution analyses for cellular response to many other external stimuli, such as intercellular signaling, the environmental pH level, mechanical forces and many others. To achieve this, the underlying population dynamic model of drug response used in PhenoPop can be replaced with another mechanistic or machine-learning derived model describing response to other stimuli. In future efforts we aim to extend PhenoPop to incorporate the role of stromal cell signaling in mediating drug-response.

The precision of PhenoPop depends on the amount of observation noise in the data. Under an exponential growth model, PhenoPop accurately performs deconvolutions on data subjected to noise with a standard deviation of up to 20% of the initial cell count, while higher noise levels lead to errors in model selection and decreased accuracy in mixture fractions and GR50 estimates. This is especially seen in the 3-population mixtures, and it is expected that the problem would be aggravated in mixtures of more than 3 populations. We note that the standard deviation-to-mean ratio reported from several of the most common automated or semi-automated cell counting techniques ranges from 1-15 %.^20,21^ At moderate noise levels in this range (standard deviation to mean ratio of 5%), PhenoPop was able to detect subpopulations as small as 1% of the total population in 2-population mixtures, while in 3-population mixtures the smallest detectable population fraction was 3 %. The precision is reduced when subpopulations have very similar GR50 values and the resolution of experimental drug concentrations does not distinguish well between them, but for predicting treatment response, distinguishing subpopulations that are almost identical is of limited clinical importance. Additionally, our study suggests that in terms of data resolution and prioritization of experimental effort, increasing the number of observed concentrations improves accuracy the most, followed by the number of time points, and then the number of replicates.

There are several limitations to the method which suggest avenues for future work. PhenoPop is currently best-suited for cancer types in which relatively large amounts of tumor material can be obtained from patients for ex vivo drug screening, such as hematological cancers. As shown in Figure 3, these drug screens can be completed in less than a week, which is an acceptable time frame for clinical decision making. Ex vivo drug screening is still challenging for solid tumours due to smaller sample size availability (typically biopsies) compared to hematological cancers. However, this field is evolving rapidly and new methods are being developed for solid tumours, such as organoid/tumoroid approaches that allow for expansion of tumor tissue and also preserve the tumor microenvironment to a certain extent. Another important consideration for the applicability of PhenoPop to solid tumors is spatial heterogeneity across different regions of a tumor, which may result in incomplete or inaccurate mixture deconvolutions. However, in situations where multiregion sampling is available, PhenoPop can be used to perform a deconvolution analysis on each sampled region, thus providing a spatially resolved profile of drug-response heterogeneity throughout the tumor.

PhenoPop currently produces a heterogeneity profile for each patient sample with respect to each treatment in a drug-screen panel. While this information is useful for identifying successful single-agent therapies and for optimizing or designing their therapeutic schedules, combination therapy design requires joint deconvolution analyses that elucidate the mapping between heterogeneity profiles for multiple drugs. This task will necessitate additional data from combination drug screens, and further methodological development in experimental design to identify tractable subsets of combination screening experiments that are necessary for identifying these joint deconvolution profiles. We plan to address this problem in future work. Another area of future work is the expansion of PhenoPop into a Bayesian framework where the incorporation of expert knowledge, genetic information or experience from previous experiments with the drug of interest could be incorporated into PhenoPop by the use of informative priors. This would enable inference on screening datasets with fewer replicates and/or time points.

Accurate, efficient techniques for profiling of heterogeneity across multiple axes are important foundations for personalized treatment decision-making. In this work we have demonstrated that PhenoPop can provide vital insights into the diversity of drug response amongst tumor cells. This framework, enabled by mixture population dynamic modeling of response to therapy, utilizes bulk drug screen data and alleviates the need for costly single-cell methods in profiling tumor heterogeneity. Although we focus here on tumor drug-response heterogeneity, the PhenoPop framework can also be applied to detect and profile heterogeneous cellular response to other stimuli, such as stromal content, nutrient/oxygen deprivation, and epigenetic modifiers. This general framework can also be applied beyond cancer to other biological settings in which reproducing populations harbor heterogeneous responses to environmental stimuli, such as the response of bacterial or viral populations to antibiotic or antiviral therapies.

## Limitations of Study

We next performed a computational study using synthetic drug screen data to identify experimental design strategies that enhance PhenoPop accuracy, and to explore the limitations of the method.

### PhenoPop experimental design recommendations

We first considered the relative importance of experimental resolution in drug concentration, time points, and replicates in PhenoPop performance. Figure 5 shows the average gain in accuracy for a mixture of 2 populations (one sensitive, one resistant) when either the replicates *R*, the number of concentrations *N_c_* or the number of time points *N_t_* are increased while the others are held constant at the value three. To compare the accuracies, 27 two-sided t-tests were made, since 3 effects (increasing *R*, *N_c_*, and *N_t_*) were compared pairwise at 3 sample sizes (5, 9 and 17), in 3 different comparison measures. To account for multiple testing, the family-wise error rate was controlled to be below 0.05 using the Bonferroni correction.

**Figure 5:**
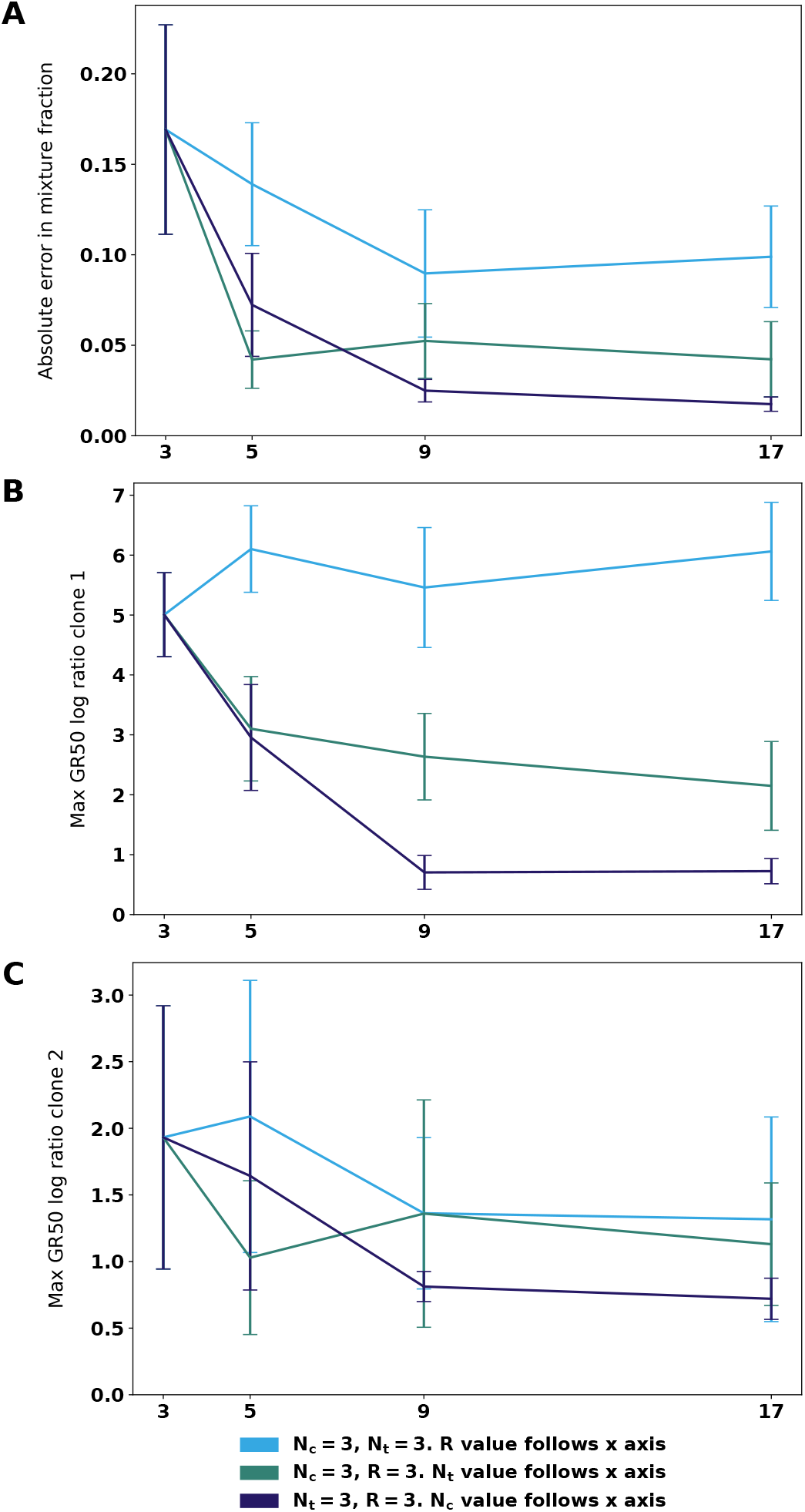
Comparison of accuracy gain in mixture fraction and GR50 values when increasing either the number of replicates (R), the number time points (*N_t_*) or the number of concentrations (*N_c_*) while keeping the other two equal to 3. The inference was carried out on 30 datasets generated from a mixture of 40% sensitive and 60% resistant cells. The standard deviation of the observation noise was equal to 10% of the initial cell count. The random seed for the noise was the only parameter varying between the 30 datasets. In A, the accuracy metric is absolute error in inferred mixture parameter; in B and C the metric is log(max(*GR50_inferred_/GR50_true_, GR50_true_/GR50_inferred_*)), chosen to address the logarithmic scale of the concentrations. The plots show mean accuracy metrics with 95% confidence intervals for the mean (t-distribution with 29 degrees of freedom). The number of subpopulations (2) was assumed known, and model selection was not performed.

We find that for accuracy in the mixture parameter, increasing the number of concentrations or time points gives significantly higher precision than increasing the number of replicates to the same amount. Similarly, to enhance accuracy in the GR50 value of the sensitive population, increasing either the number of concentrations or number of time points gives significantly higher precision compared to increasing the number of replicates by the same number. In addition, increasing the number of concentrations to 9 or 17 is significantly better than increasing the number of time points similarly. No significant differences were found for estimating the GR50 of the resistant population.

### Noise level

We also studied how increasing levels of measurement noise in the data (e.g. in cell counting) impact the precision of the deconvolution results. Results of these tests are shown in Figure 6, where the same synthetic data with increasing levels of measurement noise were used as inputs to PhenoPop. We found that for noise levels up to a standard deviation equal to 20% of the initial cell count, PhenoPop is able to correctly deconvolve the bulk response signal into the correct components. Beyond this noise level mixture fractions are off by more than 10% in 2-population mixtures, and populations may go undetected in 3-population mixtures. Supplemental figure S5 shows how model selection was performed in these cases.

**Figure 6:**
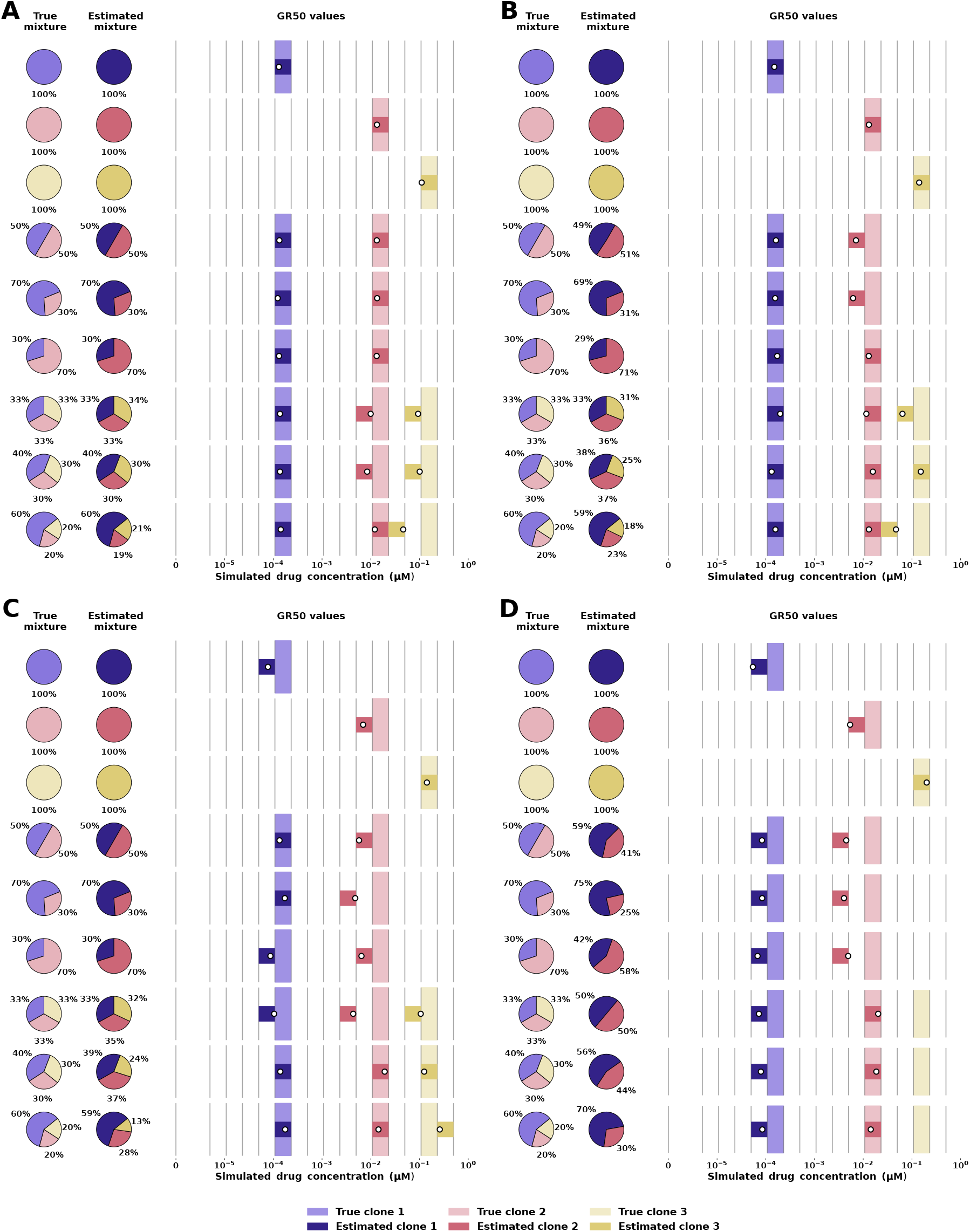
Deconvolution results for different noise levels. True and estimated mixture fractions and GR50 values for synthetic data with observation noise with standard deviation equal to A 1%, B, 10%, C 20%, D 50% of the initial cell count. Each row corresponds to an independent experiment. On the left, the true and estimated mixture percentages are shown. On the right, the grid of vertical lines corresponds to drug concentrations. The true GR50 region is marked by a light-colored bar extending between two adjacent concentrations, and the estimated GR50 region is marked by a narrower darker-colored bar. The point estimate of the GR50 is marked with a white circle. See Supplemental Figure S5 for model selection.

### Small mixture fractions

To determine how small population fractions PhenoPop is able to detect, inference was performed on simulated data with a range of small mixture fractions, with a noise level of 5% of the initial cell count. We found that in 2-population mixtures, PhenoPop was able to detect populations at frequencies as low as 1 percent. In 3-population mixtures, PhenoPop was able to detect populations with mixture fractions of 3 percent and higher. At noise level of 5%, the estimated mixture parameters were within 1% of the true value and the estimated GR50 values were always within two GR50 regions of the true value. Figure 7A shows these results. The figure also shows that it is harder to detect two small populations mixed with a large population (bottom row), than it is to infer one small population mixed with two larger ones (fifth row). Supplemental Figure S6A shows how model selection performed for these cases.

**Figure 7:**
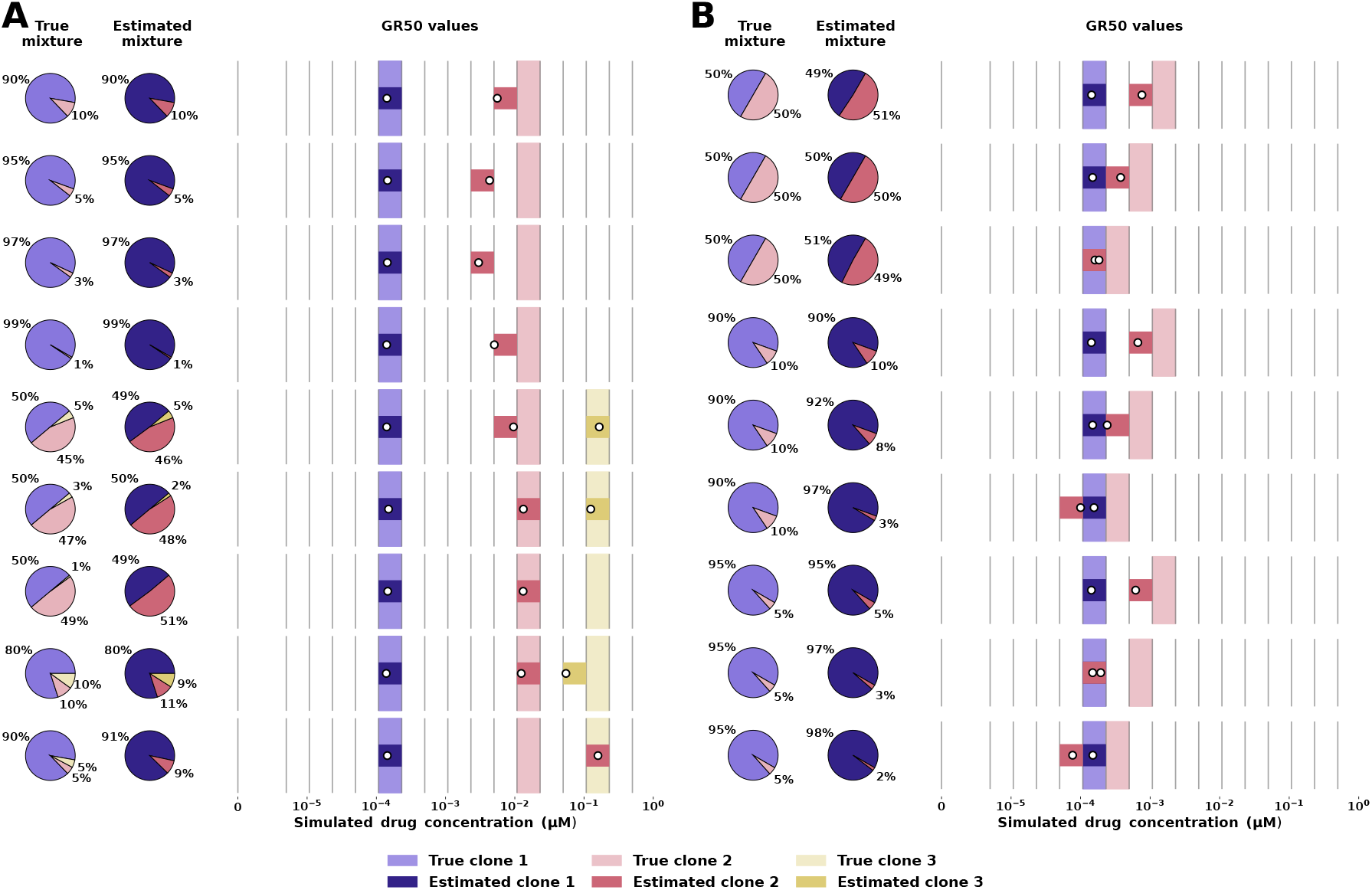
Deconvolution results for populations with small populations and close GR50 values. True and estimated mixture fractions and GR50 values for synthetic data with observation noise with standard deviation equal to 5% of the initial cell count. Each row corresponds to an independent experiment. On the left, the true and estimated mixture percentages are shown. On the right, the grid of vertical lines corresponds to drug concentrations. The true GR50 region is marked by a light-colored bar extending between two adjacent concentrations, and the estimated GR50 region is marked by a narrower darker-colored bar. The point estimate of the GR50 is marked with a white circle. See Supplemental Figure S6 for model selection.

### Subpopulation similarity

We performed computational experiments to determine the degree of similarity between component subpopulations beyond which PhenoPop was unable to detect distinct populations. We tested a set of 2 similar mixed populations, at a noise level of 5% of the initial cell count. We found that PhenoPop was able to detect populations whose GR50 values were as close as 2 GR50 regions apart. For such close populations, the estimate of the mixture parameters were within 2% of the true value and the estimated GR50 values were within 1 GR50 region of the true value, even for mixtures as unbalanced as 90:10. The results are shown in Figure 7B. The figure’s third, sixth and ninth rows show that the inferred GR50 values may overlap or swap position if the true GR50 values are less than 2 GR50 regions apart. The figure’s eighth row shows that for mixtures of 5% or smaller, the inferred GR50 values can overlap even when the true GR50 values are 2 GR50 regions apart. Figure S6B shows how model selection performed for these cases.

## Supporting information

Multiple myeloma patient information

## Acknowledgements

The numerical computations were performed on resources provided by UNINETT Sigma2 - the National Infrastructure for High Performance Computing and Data Storage in Norway. The project received funding from the UiO:LifeScience initiative through the convergence environment grant PerCaThe. A.K.L, E.M.M. and A.F. were supported by the center for research-based-innovation BigInsight under grant 237718 of Norges Forskningsråd. J.F. and K.L. were supported by the Fulbright US-Norway Foundation. J.F., K.L., and J.N. were supported by the University of Oslo-University of Minnesota Norwegian Centennial Chair Grant. J.F. was supported by the US National Science Foundation under grant number DMS-2052465. K.L. was supported by the US National Science Foundation under grant number CMMI-1552764. We acknowledge funding from the Research Council of Norway with project numbers 294916, 261936, 309273 and 262652; the Norwegian Cancer Society with project number 182524; and the Norwegian Health Authority South-East with project number 2019096. The authors also acknowledge the Centre for Digital Life Norway for supporting the partner projects PerCaThe and PINpOINT. We thank the Digital Scholarship Center, University of Oslo, for insightful advice on the visual representation and communication of our research findings.

## Author Contributions

Conceptualization, A.K.L., K.L., A.F., and J.F. Formal analysis, A.K.L., E.M.M., J.N., E.H., P.A., K.L., A.F., and J.F. Funding acquisition, A.K.L., K.L., A.F., and J.F. Investigation, E.M.M., D.S.T., M.G., S.M.M., F.S., K.T., and J.M.E. Methodology, A.K.L., E.M.M., K.L., A.F., and J.F. Resources, S.M.M., F.S., K.T., J.M.E., and A.F. Software, A.K.L., E.M.M., L.S., J.N., E.H., and P.A. Validation, A.K.L., E.M.M., D.S.T., M.G., L.S., S.M.M., F.S., K.T., and J.M.E. Visualization, A.K.L., E.M.M., M.G., K.L., and J.F. Writing – original draft, A.K.L., E.M.M., K.L., A.F., and J.F. writing – review & editing, A.K.L., E.M.M., S.M.M., F.S., K.T., J.M.E., K.L., A.F., and J.F.

## STAR Methods

### Key resources table

#### Resource availability

##### Lead contact

Requests for further information should be directed to and will be fulfilled by the lead contact, Jasmine Foo.

##### Materials availability

This study did not generate new unique reagents.

##### Data and code availability

- All data and code used in this article are publicly available in the Github page of Oslo Center for Biostatistics and Epidemiology. The Phenopop repository contains the data and code needed to reproduce the results in this article, and a user-friendly example of how to run PhenoPop in Matlab. Additionally, the pyPhenoPop repository contains a user-friendly example of how to run PhenoPop in Python, and can be easily installed as a package through pip. The repositories are also backed up at Zenodo, for PhenoPop and pyPhenoPop separately. DOIs are listed in the key resources table. Any additional information required to reanalyze the data reported in this paper is available from the lead contact upon request.
- All original code has been deposited at Zenodo and is publicly available as of the date of publication. DOIs are listed in the key resources table.
- Any additional information required to reanalyze the data reported in this paper is available from the lead contact upon request.

#### Experimental model and subject details

##### Ba/F3 cell line experiments

###### Preparation of cell lines

BCR-Abl-T315I expressing plasmid was established by site-directed mutagenesis of p210 BCR-Abl (Addgene 27481) using QuickChange II XL (Agilent Technologies) with the forward primer 5’ GGGAGCCCCCGTTCTATATCATCATTGAGTTCATGACCTACG 3’ and the reverse primer 5’ CGTAGGTCATGAACTCAATGATGATATAGAACGGGGGCT CCC 3’ for T315I. To generate cells stably expressing BCR-Abl (imatinib-sensitive) and BCR-Abl-T315I (imatinib-resistant), parental Ba/F3 cells were transfected with the appropriate plasmids by electroporation using Amaxa biosystems nucleofecor II and stable cells were established by selecting with medium containing 500 μg/mL Geneticin (Gibco, UK) and lacking the growth factor IL3 (BCR-ABL activity can overcome the requirement for IL3 of untransformed parental cells for survival/proliferation).^22^ Furthermore, Ba/F3 cells expressing BCR-Abl were stably transfected with GFP expression, pRNAT-H1.1/Hygro plasmid from Genscript (Piscataway NJ, USA). The resulting subpopulations exhibited distinctive phenotypic differences upon treatment with Imatinib.

###### Cell cultures

Parental Ba/F3 cells were maintained in RPMI-1640 supplemented with 10% heat-inactivated Fetal Bovine Serum (FBS), 7.5 ng/mL IL3 and 1% penicillin and streptomycin at 37°C under a humidified atmosphere containing 5% CO2. Ba/F3 cells stably expressing BCR-Abl and BCR-Abl-T315I were maintained in medium lacking IL3.

###### Experimental procedures

Cells were harvested at 70-80% confluence, stained with trypan blue (ThermoFisher, UK), and counted with a Countess 3 Automated Cell Counter (Life Technologies). Mono- and co-cultures were seeded at different initial ratios in 384 well microplates (Greiner Bio-One) that contained different concentrations of imatinib (Cayman, USA). Imatinib ranging from (0 - 5 μM) was dispensed using an Echo acoustic liquid dispenser (Labcyte, San Jose, CA, USA) in seven replicates per condition. Then time-lapse microscopy images were obtained for bright field and GFP using IncuCyte (Essen BioScience, UK) every 3 hours over the course of 72 hours.

###### Image processing

Images were processed with the open-source software ImageJ.^33^ Images were background subtracted, converted to 8-bit, bandpass filtered, sharpened, contrast enhanced, and thresholded. Then images were converted to binary images, watershed segmentation was performed, and raw cell numbers were extracted.

##### NSCLC cell line experiments

###### Cell cultures

HCC827 and H1975 cell lines were maintained in RPMI-1640 media supplemented with 10% Fetal Bovine Serum and 1% penicillin and streptomycin under standard cell culture growth conditions (37°C and 5% CO2).

###### Experimental growth assay

Tumor cells were seeded in 96-well black walled plates at 5,000 cells per well. The following day, the cells were treated with erlotinib at various concentrations (0, 0.1, 1, 10uM). Cell counts were determined at 0, 24, and 48 hours post drug treatment using the Cellomics Arrayscan High Content Screening Platform. Briefly, cells were stained with 5 μg*/*mL Hoechst 33342 (nuclear marker to determine total cell count) and 5 μg*/*mL Propidium Iodide (PI - vital dye to determine dead cells) for 45 minutes prior to imaging. The average intensity for Hoechst and PI was determined for each cell to classify as live or dead. Each condition was performed in replicates of four. For admixture experiments, each cell line was labeled with a different CellTracker dye (CellTracker orange CMTMR and H1975 labeled with CellTracker green CMFDA). The cells were mixed at the specified ratios (total 5,000 cells/well) and imaged following the procedures outlined above.

##### Drug screen of Multiple Myeloma patient samples

###### Patient samples

The multiple myeloma (MM) patients enrolled in this study were recruited from the Oslo Myeloma Center at Ullevål Oslo University Hospital under the Regional Committee approval for Medical and Health Research Ethics of South-Eastern Norway (REC-2016/947). The MM samples were obtained following written informed consent in compliance with the Declaration of Helsinki.

###### Primary MM cells processing

Bone marrow samples from 5 relapsed myeloma patients were collected in ACD tubes. Details about patient ID, treatment lines and refractory status are provided in Supplemental Table 1. A Lymphoprep TM (Stemcell Technologies) density gradient centrifugation method was used to obtain bone marrow mononuclear cells (BMMCs) from patient samples. As described in Wang et al.,^34^ after CD8 T cell depletion by Dynabeads (Life Technologies), BMMCs were then stimulated by activated T helper cells in the presence of Human T-activator CD3/CD28 Dynabeads (Life Technologies) and 100U/mL human interleukin-2 (hIL-2, Roche, Mannheim, Germany). After 24h, CD138+ MM cell enrichment was performed from the BMMC fraction by immune-magnetic microbeads CD138+ (Milteny Biotec, Bergisch Gladbach, Germany).

###### Drug sensitivity assay

CD138+ MM cells (200,000 cells/mL) derived from activation assays were treated with drugs at 9 concentrations using a drug customized concentration range (0,1-10,000), as described in Giliberto et al.^26^ The drug panel included clinically relevant anti-myeloma drugs, Dexamethasone (0,1-10,000), ixazomib (1-10,000), thalidomide (0,1-10,000), selinexor (0,1-1000), melflufen (0,1-1000) and venetoclax (0,1-10,000). Pre-printed drug plates were made by an acoustic dispenser (Echo 550, LabCyte Inc., San Jose, CA, USA), by the Chemical Biology Platform, NCMM, University of Oslo. Control agents included a negative control, 0,1% solvent solution dimethyl sulfoxide (DMSO), and a positive control 100 uM benzethonium chloride (BzCl). In brief, MM cells were diluted in culture medium (RPMI 1640 medium supplemented with 10% fetal bovine serum, 2mM L-glutamine, penicillin (100U/mL), streptomycin (100 μg/mL), and 25 μL of cell suspension was transferred to 384-well plates using a Certus Flex liquid dispenser (Fritz Gyger, Switzerland). Afterward, plates were incubated at 37°C and 5% CO2 humidified environment. Cell viability was measured at 4 different time points (0h-96h), using the CellTiterGlo (Promega, Madison, WI, USA) ATP assay according to manufacturer’s instructions and with an Envision Xcite plate reader (Perkin Elmer, Shelton, CT, USA) to measure luminescence.

### Method details

Given a set of experimental drug-screen data on a bulk tumor sample, PhenoPop solves a series of optimization problems to identify individual subpopulations within the sample and to estimate their frequencies and drug sensitivities. This problem is challenging because it requires simultaneous estimation of the number of individual subpopulations present, their frequencies in the population and their drug response characteristics, all based on noisy observations of the total cell population. Our solution to this problem is enabled by the introduction of a mixture population dynamic model of the tumor in which the growth rate dependence on drug concentration follows a Hill-type functional form (see equation (3)).

#### Model of dose-dependent population dynamics

PhenoPop relies upon an underlying model of heterogeneous tumor population dynamics *in vitro*. The growth of a single population of cells with homogeneous drug response is modelled by

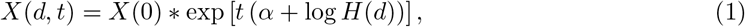

where *X*(*d,t*) is the number of cells at time *t* under drug concentration *d, X*(0) is the initial population size, *α* is the intrinsic growth rate of the population in the absence of drug, and *H*(*d*) is a classic sigmoidal function describing the dependence of the population growth rate on drug concentration *d*:

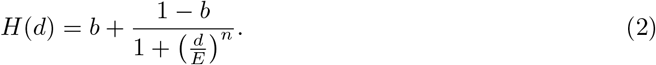

The parameters of this function control the shape of the sigmoid: *b* ≥ 0 reflects the maximum effect of the drug, *E* is the log concentration at which 50 percent of the maximum effect is achieved, and *n* > 0 controls the steepness of the response. This form of the growth rate, *r*(*d*) ≡ *α* + log *H*(*d*), is chosen so that the predicted cell viability curve, which is the treated viable cell population size normalized by the untreated viable cell population size at a fixed time, exhibits the standard Hill-shaped dependence on drug concentration that is empirically observed in viability assays.^16^ Supplemental Figure S1 demonstrates that this model accurately recapitulates experimental cell viability dependence on drug concentration in two BCR-ABL positive Ba/F3 cell lines (with and without the T315I mutation) treated with the tyrosine kinase inhibitor imatinib. Note that since we are studying *in vitro* populations prior to confluence, an exponential growth model is appropriate.

To extend the monoclonal growth model in equation 1 to a population composed of several subpopulations, each with a specific own drug response dynamics, we denote the growth parameters of the i-th subpopulation by *α_i_*, *b_i_*, *E_i_*, *n_i_*. Then the model of a cell population with *S* subpopulations under drug concentration *d* at time *t* is:

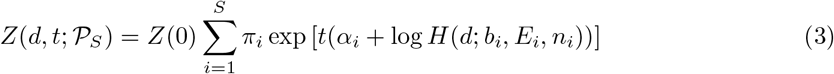

where *Z*(0) is the total initial population and *π_i_* is the initial mixture fraction of the *i*-th subpopulation 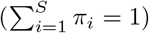. Here, 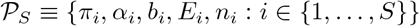 denotes the set of parameters for S populations, and the parameters of the Hill function *H*(*d*; *b_i_*, *E_i_*, *n_i_*) are written explicitly to emphasize the individual drug response profile of each subpopulation. Under this formulation, we need to estimate, on the basis of the drug screen data, the unknown number of subpopulations *S* and the corresponding parameters 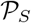. Note that in this case, the heterogeneous population is modelled as a mixture of populations in which individual subpopulations are assumed to grow independently. In the Supplemental Information, we consider a case in which interaction between subpopulations is incorporated.

#### Estimation procedure

As input, PhenoPop takes bulk tumor sample drug screening observations, in the form of total cell counts at a series of time points and drug concentrations. A variety of experimental techniques is commonly used to generate such observations of cell population counts in drug screening. For example, tetrazolium reduction assays (e.g. MTT, MTS), protease viability markers (e.g. GF-AFC), ATP assays (e.g. Cell Titer-Glo), and more recently developed real-time assays (e.g. Real-Time Glo, live-cell imaging).^35,36^ The PhenoPop methodology is capable of using experimental input from any of these assays, as long the measurements provide viable cell count or a proxy quantity (e.g. fluorescence intensity) that is proportional to the cell number. Generally, real-time techniques may yield superior deconvolution results due to a reduction in the total noise of the data set.

Given a set of experimental drug-response data on a bulk tumor sample, PhenoPop solves a series of optimization problems to deconvolve and characterize individual subcomponents of the bulk sample in terms of varying drug sensitivity profiles. In particular, each experimental observation, denoted by 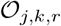, corresponds to a cell population number measured under drug concentration *d*(*j*) where *j* ∈ {1,…, *C*}, time point *t_k_*, where *k* ∈ {1,…, *T*}, and replicate *r* ∈ {1,…, *R*}. We denote the total set of observations by 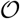.

For simplicity, we will first assume that there are *S* subpopulations. Our statistical model of experimental observations will be based on the deterministic model in equation 3. In particular, we model each experimental observation as an independent standard Gaussian random variable with mean 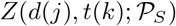) and standard deviation *σ*(*d*(*j*), *t*(*k*)). Note that we allow the standard deviation *σ* to vary with dose and time. This is because at low doses and high times we expect a larger variance due to the larger cell counts. Therefore we define

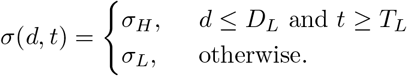

Our standard deviation is thus characterized by four parameters, *σ* = (*σ_L_*, *σ_H_*, *D_L_*, *T_L_*). We will denote the set of time-dose observations where we use standard deviation *σ_H_* by *I_H_*, and the set where we use *σ_L_* by *I_L_*. We denote their cardinalities by |*I_H_*| and |*I_L_*|.

Assuming *S* subpopulations we can use this model to write the log-likelihood as

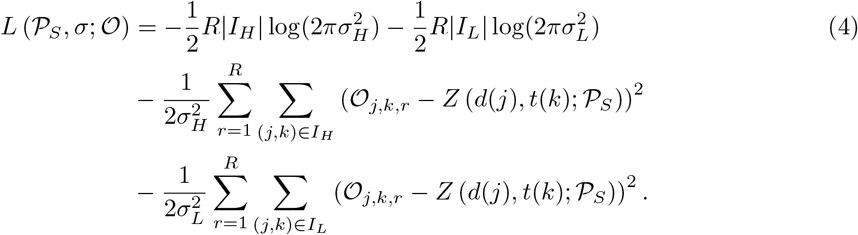

For a fixed *S*, we thus compute the maximum likelihood estimates of the model parameters by solving the optimization problem

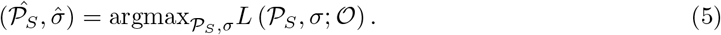

##### Model selection using the elbow method

We observed that traditional model selection criteria like BIC, AIC and the likelihood-ratio test do not consistently select the correct model in simulated cases and cell cultures of known admixtures, tending instead to select larger numbers of subpopulations. This tendency of the AIC and BIC to overestimate the number of components in a mixture model has been documented in other contexts (e.g. Celeux et al.^37^) though the reasons remain unclear. Instead, a heuristic known as the elbow method was used. To infer the number of subpopulations in the mixture, PhenoPop is fitted to the data repeatedly, for each number of subpopulations *S* in *S* = {1, 2,…, *S*_max_} in turn, and the *S* negative log-likelihood values are recorded. We then plot the negative log-likelihood values as a decreasing function of *S*, and observe the number of subpopulations corresponding to which the negative log-likelihood does not decrease significantly further. This means that no useful increase in model accuracy is gained by including another additional population. This point of inflection of the negative log-likelihood is called the elbow of the curve. The optimal number of populations is then chosen by the experimenter through visual inspection. The resulting estimate 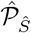 contains the inferred population’s drug response substructure: the estimated number of populations along with the estimated mixture frequency and estimated drug sensitivity *GR*_50_ of each subpopulation. This method is known as the elbow method, and it is a well-known heuristic for model selection in cases where the model fit generally increases with complexity. Model selection for all experiments is shown in Supplemental Figures S3, S4, S5, and S6. Note that the negative log-likelihood value at the true global minimum of the negative log-likelihood should in theory decrease monotonically as the number of subpopulations increases, since extraneous mixture parameters can always be set to zero. However, as the number of subpopulations increases the complexity of the optimization problem also increases, so in practice negative log-likelihood values may become non-monotonic due to the difficulty of obtaining convergence to the exact global minimum within the available iterations.

### Quantification and statistical analysis

#### Optimization methodology

The maximum likelihood estimate of the parameters 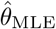 was obtained by maximizing the loglikelihood in equation 4, subject to constraints that were placed on the range of each parameter. This constrained optimization problem was performed using the function fmincon from the MATLAB Optimization Toolbox in Matlab version R2020b,^38^ with the default interior-point optimization method. To combat converging to suboptimal local minima, the log-likelihood was maximized repeatedly and independently, by starting from N_optim_ different random initial positions for the parameter *θ*, sampled uniformly within their allowed range (except for the parameter *E*, which was sampled log-uniformly within the bounds). Among the N_optim_ minima, the one with the highest log-likelihood value was chosen as estimate 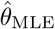.

#### Calculation of *GR*_50_ values

The viability curve and associated metrics of drug response (e.g. *IC*_50_, *EC*_50_) typically exhibit dependence on the timing of data collection.^19^ We form a growth rate curve by inferring the growth rate *r*(*d*) at each tested dose level *d*. In contrast to the viability curve the growth rate curve does not have a hidden dependence on the duration of the experiment, assuming exponential growth. Once the parameters of the model in equation 3 are estimated for each subpopulation using the inferential procedures above, the *GR*_50_ for each subpopulation can be explicitly determined using the set of parameters (*α_i_*, *b_i_*, *E_i_*, and *n_i_*). Following Sorger et al.,^19^ we characterize dose-response of clones with a *GR*_50_ value. This number represents the dose at which the cellular growth rate experiences half of its total reduction. In particular, suppose that we are interested in a homogeneous population with the growth rate at dose *d* given by

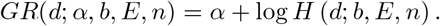

Note that we will generally suppress the dependence on parameters and simply write *GR*(*d*). If the maximum dose administered is *d_m_*, and the minimum dose administered is 0, then the median growth rate is *r_m_* = (*GR*(0) + *GR*(*d_m_*))/2. We then define the *GR*_50_ as the dosage that results in this growth rate, i.e., the value d such that *GR*(*d*) = *r_m_*. We can then solve to obtain

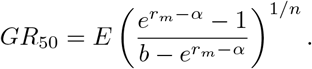

#### Generation of synthetic population data

By defining a number of populations S and a parameter set 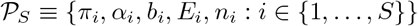, synthetic data can be generated in a deterministic manner with equation 3. Table 1 shows the parameters {*α_i_*, *b_i_*, *E_i_*, *n_i_*: *i* ∈ {1, 2, 3}} of the blue, red and yellow populations in Figures 2, 6, and 7.

**Table 1:** Parameters for simulated data.

| | $\alpha$ | b | E | n |
| --- | --- | --- | --- | --- |
| Clone 1 (blue) | 0.03 | 0.3 | 0.0001 | 3.0 |
| Clone 2 (red) | 0.03 | 0.4 | 0.01 | 3.0 |
| Clone 3 (yellow) | 0.03 | 0.5 | 0.1 | 3.0 |

For the synthetic validation, simulated data with initial population size of *Z*_0_ = 1000 cells were generated for the following 9 mixtures of the three cell populations in Table 1: [1,0,0], [0,1,0], [0,0,1], [0.5,0.5,0], [0.7, 0.3, 0], [0.3, 0.7, 0], [1/3, 1/3, 1/3], [0.4, 0.3, 0.3] and [0.6, 0.2, 0.2].

We chose 17 simulated drug concentrations. One equal to zero, the rest spaced log-linearly in a region designed to cover the GR50 values of the simulated populations. The simulated concentrations were: [0, 0.00000500, 0.0000108, 0.0000232, 0.0000500, 0.000108, 0.000232, 0.000500, 0.00108, 0.00232, 0.00500, 0.0108, 0.0232, 0.0500, 0.108, 0.232, 0.5] μM. Cell counts were measured at 12-hour intervals from 0 to 96 hours, and 4 replicates of the simulation were carried out, where the only difference between the replicates was the randomly sampled observation noise: A random noise term was added to each observed cell count, sampled from an independent and identically distributed (i.i.d.) Gaussian distribution with mean 0 and standard deviation ranging from 1 to 50% of the initial cell count. Any negative cell count caused by the additive noise was set to zero. This gives the following expression for the generated observation 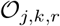 with concentration number *j* at time *k* for replicate *r*:

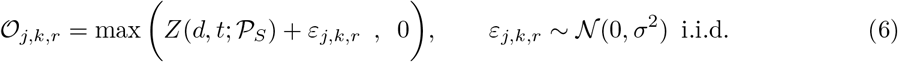

#### Model of multiple myeloma under treatment

Inspired by Tang et al.,^39^ we present a mathematical model of M-protein levels of a multiple myeloma patient under treatment with an anti-cancer drug. This model assumes that the patient has two subpopulations of cancer cells with distinct responses to the drug. In particular the cancer cells and M-protein levels are governed by the following system of ordinary differential equations

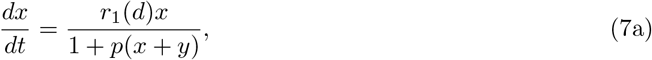

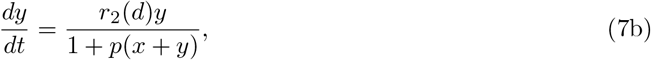

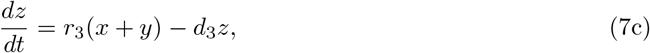

where *x* and *y* denote number of myeloma cells in subpopulations 1 and 2 respectively, and *z* denotes M-protein concentration in plasma. Parameters *r*_1_ and *r*_2_ are the net growth rates under treatment of subpopulations 1 and 2 respectively. We assume the net growth rates can be computed as

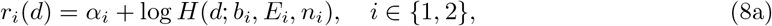

where (*α_i_*, *b_i_*, *E_i_*, *n_i_*) are the estimated parameters of subpopulation *i* using PhenoPop. The term (1+ *p*(*x* + *y*))^-1^ in equations 7a and 7b alters the growth rate of both subpopulations when the total number of cells increases. Parameters *r*_3_ and *d*_3_ are the production and decay rate of the M-protein, respectively. Inspired by Tang et al.,^39^ we use *p* = 10^-13^, *r*_3_ = 0.07 * 10^-13^ and *d*_3_ = 0.07.

#### Model parameter ranges

For model with *S* = {1, 2, 3, 4} populations, the log-likelihood was maximized *N*_optim_ = 1000 times or more to combat local minima. For each maximization, the initial estimate was sampled from within the bounds on the parameter range, which were set to the values listed below for the different datasets.

The parameter ranges for the different settings were largely similar. Some differences occur due to different concentration scales in the different experiments or due to parameter estimates hitting the boundary of the domain, in which case the range was expanded. When working with the Ba/F3 cells we needed to adjust the lower bound on the parameter *b*. Due to the complexity of the optimization problem, the solver had a tendency to push *b* towards an unrealistically low value. To address this issue we used previous observations and derived a realistic lower bound on *b*. Denote the net growth rate of the cells by *λ* = *β* — *μ*, where *β* is the birth rate and *μ* the death rate. From Milo et al.,^40^ we know that *β* ≤ .06. We can thus write *μ* = *β* — *λ* ≤ .06 — *λ_min_* = *d*_0_, where *λ_min_* is the minimum observed growth rate amongst all Ba/F3 cell line experiments. Thus the maximal possible death rate is *d*_0_, and the minimal possible net growth rate is —*d*_0_. Next note that according to our growth rate model, as the dose *d* goes to infinity the growth rate decreases to the lower limit *α* + log(*b*). Therefore we know that *α* + log(*b*) ≥ —*d*_0_. We again use that *α* ≤ .06, and based on observed data we set *λ_min_* = .04 and get *d*_0_ = 0.2. However to account for any possible errors in the method we increase *d*_0_ to be 0.07. This then gives us the lower bound log(*b*) ≥ —0.08 or equivalently *b* ≥ 0.878.

##### NSCLC data

*π_i_* ∈ [0, 1] with the inequality constraint 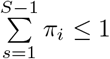
*α_i_* ∈ [0, 1] *hours*^-1^
*b* ∈ [0, 1] *hours*^-1^
*E* ∈ [0, 50] μM
*n* ∈ [0, 50]
*σ_L_*, *σ_H_* ∈ [0, 5500]

##### Ba/F3 data

*π_i_* ∈ [0,1] with the inequality constraint 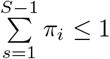
*α_i_* ∈ [0, 0.06] *hours*^-1^
*b* ∈ [0.878,1] *hours*^-1^, see comment below.
*E* ∈ [0, 50] μM
*n* ∈ [0.001, 20]
*σ_L_*, *σ_H_* ∈ [0, 2500]

##### Synthetic data

*π_i_* ∈ [0,1] with the inequality constraint 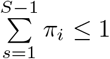
*α_i_* ∈ [0, 0.1] *hours*^-1^
*b_i_* ∈ [0.27, 1] *hours*^-1^
*E_i_* ∈ [10^-6^, 0.5] μM
*n_i_* ∈ [0.01, 10]
*S* ∈ [0, 4]
*σ_L_*, *σ_H_* ∈ [10^-6^, 5000]

##### Multiple Myeloma data

For the multiple myeloma patient data, an inital parameter range was chosen for all patients. Then if one or more of the inferred parameters happened to lie on or near the upper or lower bound, the parameter range was increased for that patient until the estimate was no longer on the bound. Therefore, the parameter for the *E* and *σ* variables are different for some of the patients.

*π_i_* ∈ [0, 1] with the inequality constraint 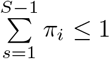
*α_i_* ∈ [-0.1, 0.1] *hours*^-1^
*b_i_* ∈ [0, 1] *hours*^-1^
*n_i_* ∈ [0.01,10]
*S* ∈ [0, 5]

The *E* parameter ranges were:

*E_i_* ∈ [10^-6^, 2] μM for patient MM2108.
*E_i_* ∈ [10^-6^, 50] μM for patient MM720.
*E_i_* ∈ [10^-6^, 5] μM for patient MM195.
*E_i_* ∈ [10^-6^, 5] μM for patient MM36.
*E_i_* ∈ [10^-6^,100] μM for patient MM1420.

The *σ* parameter ranges were:

*σ_L_*, *σ_H_* ∈ [10^-6^, 50,000] for patient MM2108.
*σ_L_*, *σ_H_* ∈ [10^-6^,1,000,000] for patient MM720.
*σ_L_*, *σ_H_* ∈ [10^-6^,150,000] for patient MM195.
*σ_L_*, *σ_H_* ∈ [10^-6^, 250,000] for patient MM36.
*σ_L_*, *σ_H_* ∈ [10^-6^,150,000] for patient MM1420.

#### Model extension to interacting populations

Our model currently ignores potential interactions between subpopulations. Based on the sample size of our current data sets we were not able to fit a more complex model that allows for interacting populations. For the situation when sufficient data are available, we propose the model below that allows for interactions between the subpopulation. Assuming that there are *S* subpopulations, for each *i* ∈ {1, …, *S*} define the function

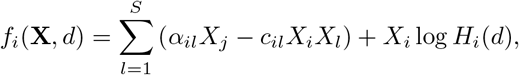

where *d* is possible drug dose and 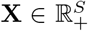. The parameter *α_il_* represent the rate at which type-*l* cells produce type-*i* cells, and *α_ii_* is the net growth rate of the type-*i* cells. We assume that each *α_il_* term is non-negative. The term *c_il_* represents the effect of population *l* on population *i*. If *c_il_* > 0 then population *l* inhibits population *i*, if *c_il_* < 0 then population *l* encourages population i to grow, and finally if *c_il_* = 0 then population *l* has no direct effect on population *i*. Note that the term *c_ii_* represents the effect of type-*i* cells on itself and we assume that *c_ii_* > 0. The parameters *α_il_* allow for inter-conversion between cell types, and the parameters *c_il_* allow for inhibition or promotion between cell types.

For dose *d*, and initial population vector 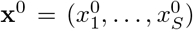, define {**X**(*t*, *d*; **x**^0^); *t* ≥ 0} as the solution to the differential equation

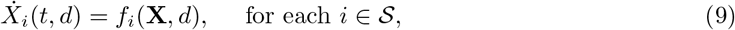

with initial condition 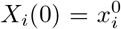. Define 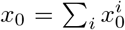 and write 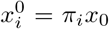. We assume that *x*_0_ is a known quantity, but the proportions {*π_i_*}_*i∈S*_ are unknown. We denote the model-predicted total population at time *t* under dose d by *X*(*t,d*). Recall that the total population is the observable variable in our model.

In this interacting population model, we have more model parameters, namely the parameter set

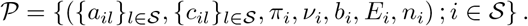

To make clear the dependence on the parameter set 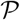, we denote the predicted total population at time *t* using *d* units of drug with parameter set 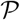 by 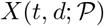.

Similar to our main model, we will start by simply using additive Gaussian noise for our measurement error. In particular, we assume that observation at dose *d_j_* and time *t_k_* is given by

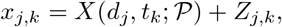

for i.i.d *N*(0, *σ*^2^) random variables *Z_j,k_*. We can then implement the same maximum likelihood estimation procedure as for our original model. This will be a more computationally challenging problem because evaluating the likelihood function will require numerically solving the non-linear differential equation 9. In addition, this inference problem is more difficult because we have a higher dimensional parameter space to search over.

## Supplemental excel tables

Supplemental table 1: Patient clinical data and drugs tested in screen assay, related to STAR Methods.

## Supplemental figures

**Figure S1:**
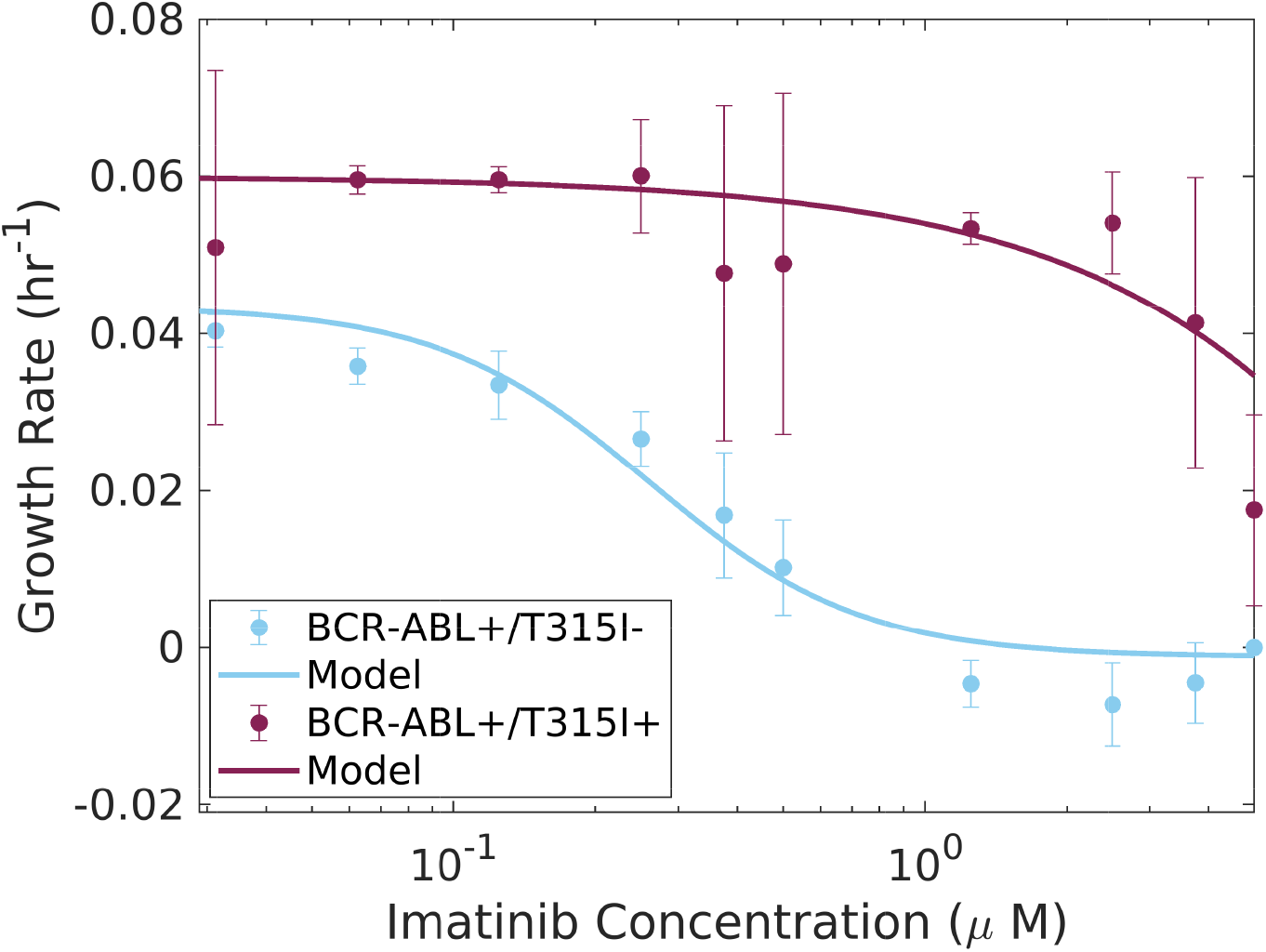
Related to STAR Methods. Model from Equation (1) of the STAR methods section accurately recapitulates experimental cell viability dependence on drug concentration in two example BCR-ABL positive Ba/F3 cell lines (with and without the T315I mutation) treated with tyrosine kinase inhibitor imatinib.

**Figure S2:**
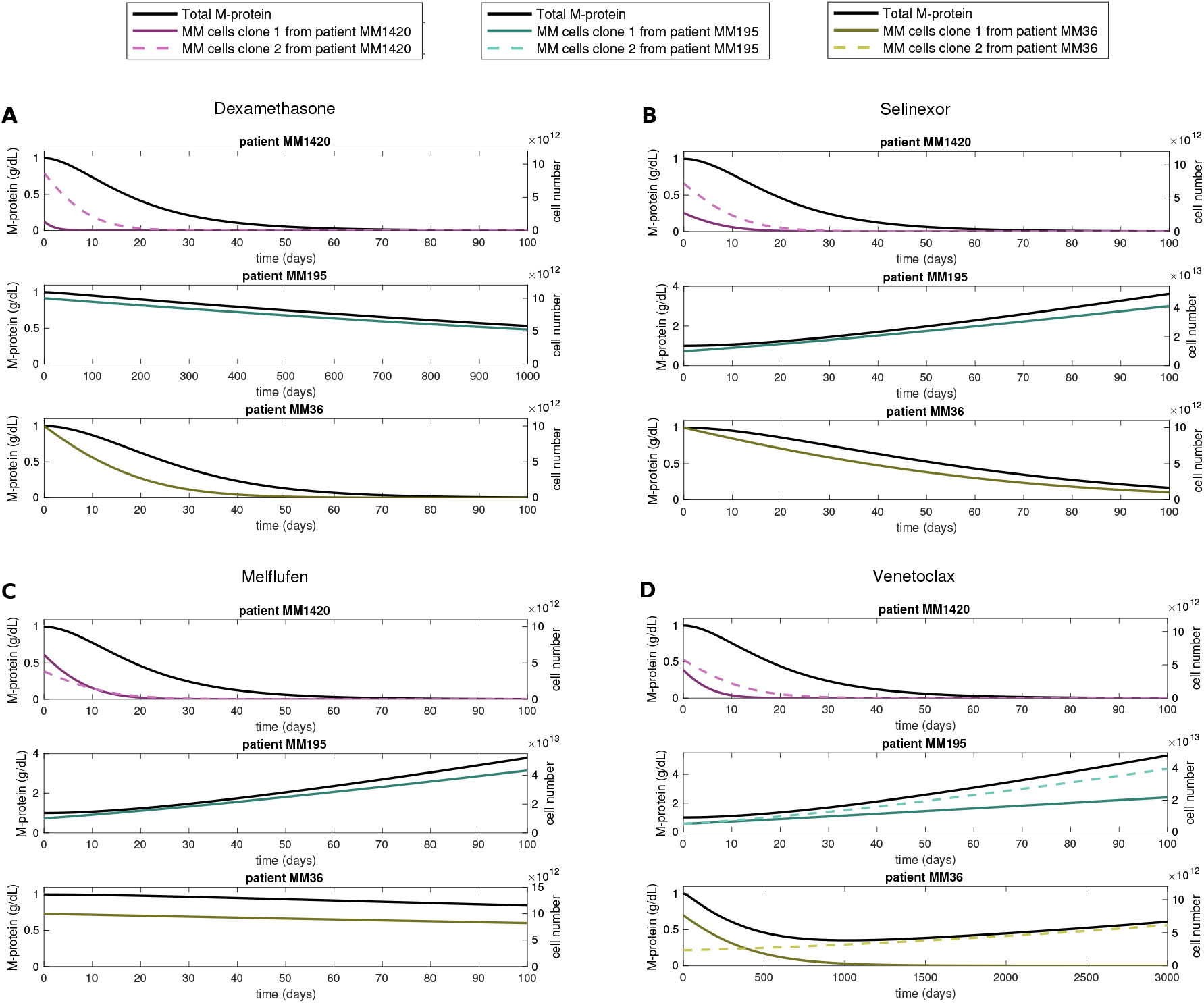
Related to Figure 4. Proof-of-concept modeling of Multiple Myeloma disease dynamics for three patients under treatment with A, Dexamethasone at 5 μM, B, Selinexor at 0.5 μM, C, Melflufen at 0.5 μM, D, Venetoclax at 2 μM, using PhenoPop deconvolution results. The estimated mixture and drug-response parameters obtained by PhenoPop (see Figure 3E) define the initial percentage of cells and drug-response for each clone and patient. Cells from both clones are assumed to produce monoclonal protein (M-protein), which can be used as a proxy for tumor burden. For easier comparison, we assume that all three patients start with a total of 10^12^ abnormal plasma cells (cell number shown in the right y-axes) and 1*g*/*dL* M-protein (shown in the left y-axes). See the STAR methods section for a description of the mathematical model.

**Figure S3:**
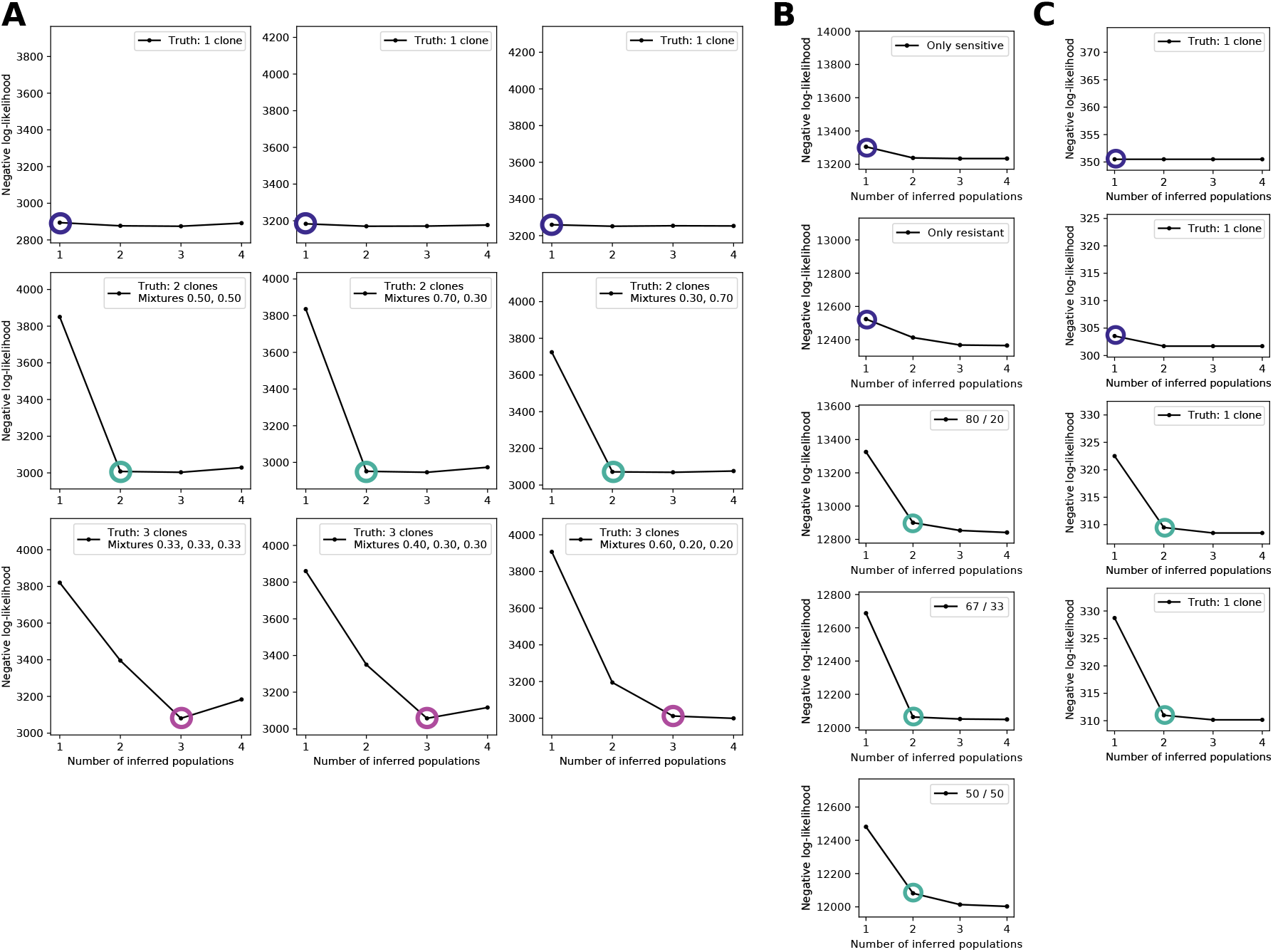
Related to Figure 2. Elbow plots showing the negative log-likelihood values for panel A, B and C in Figure 2 respectively, with the selected model marked by a circle. The color of the circle also indicates the selected model: blue for 1 population, teal for 2, dark magenta for 3.

**Figure S4:**
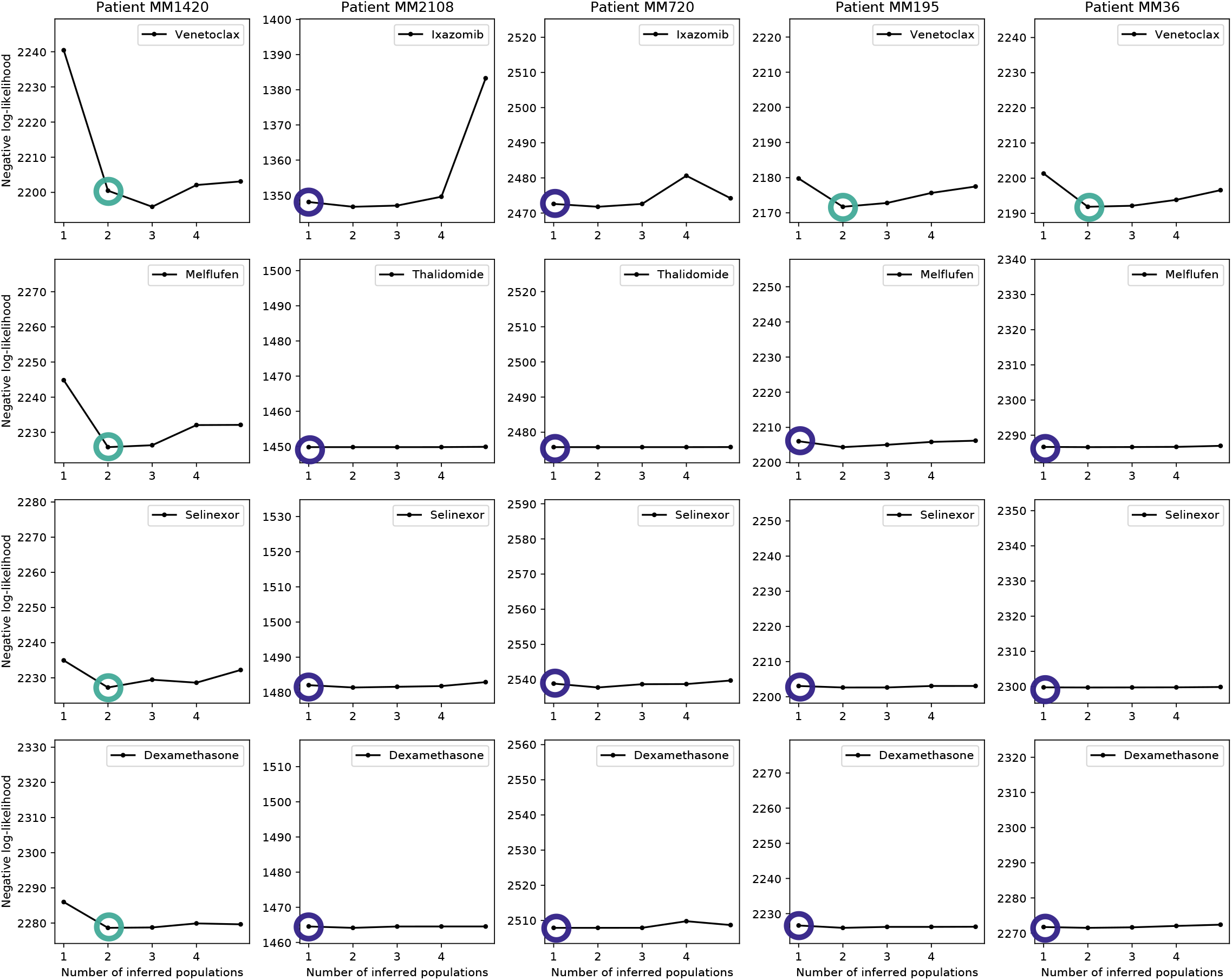
Related to Figure 3. Elbow plots showing the negative log-likelihood for all drugs for all multiple myeloma patients, with the selected model marked by a circle. The color of the circle also indicates the selected model: blue for 1 population, teal for 2, dark magenta for 3.

**Figure S5:**
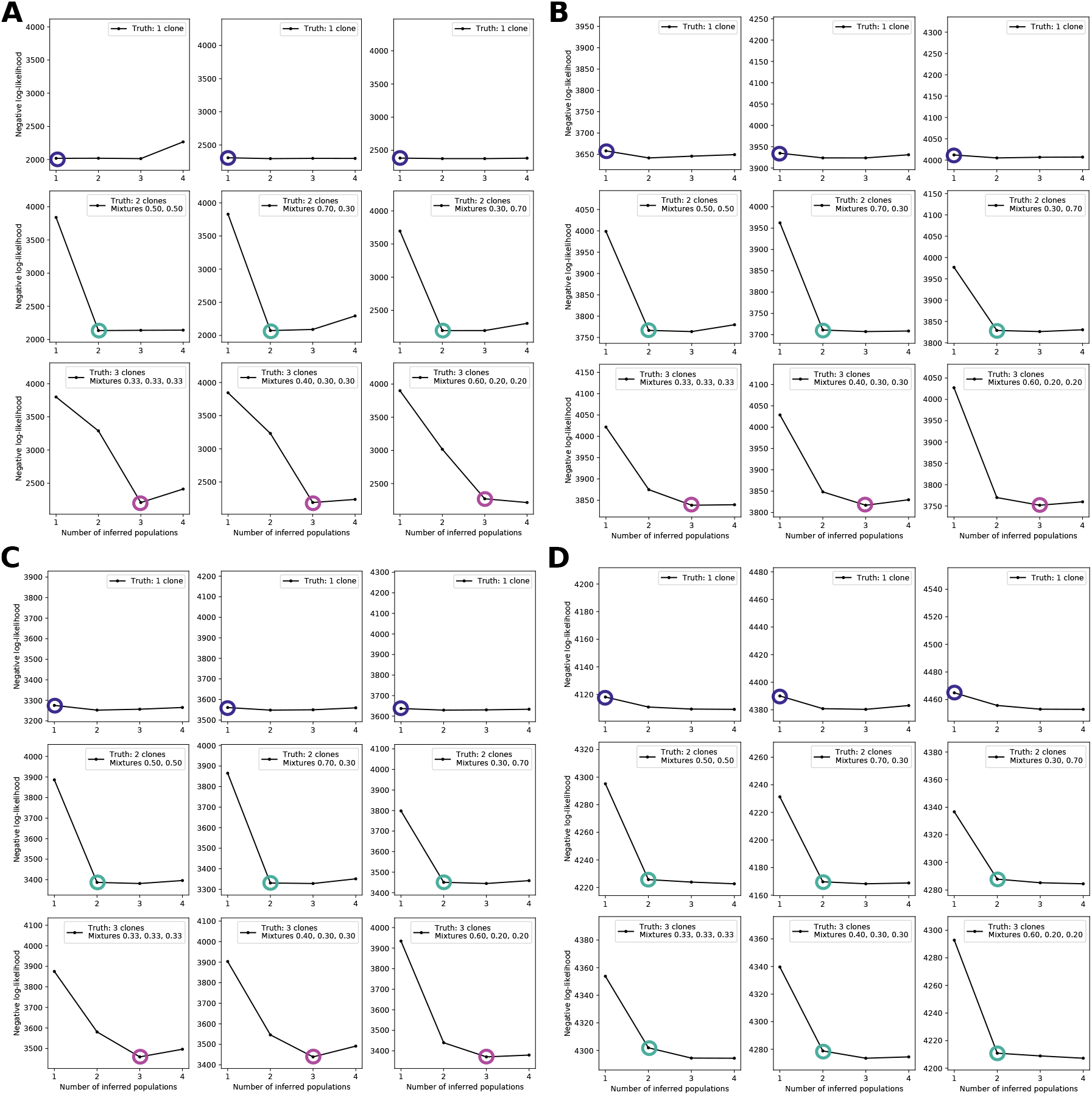
Related to Figure 6. Elbow plots showing the negative log-likelihood for all cases in figure Figure 6; simulated data with observation noise with standard deviation equal to A 1%, B, 10%, C 20%, D 50% of the initial cell count, respectively. The selected model is marked by a circle, and the color of the circle also indicates the selected model: blue for 1 population, teal for 2, dark magenta for 3.

**Figure S6:**
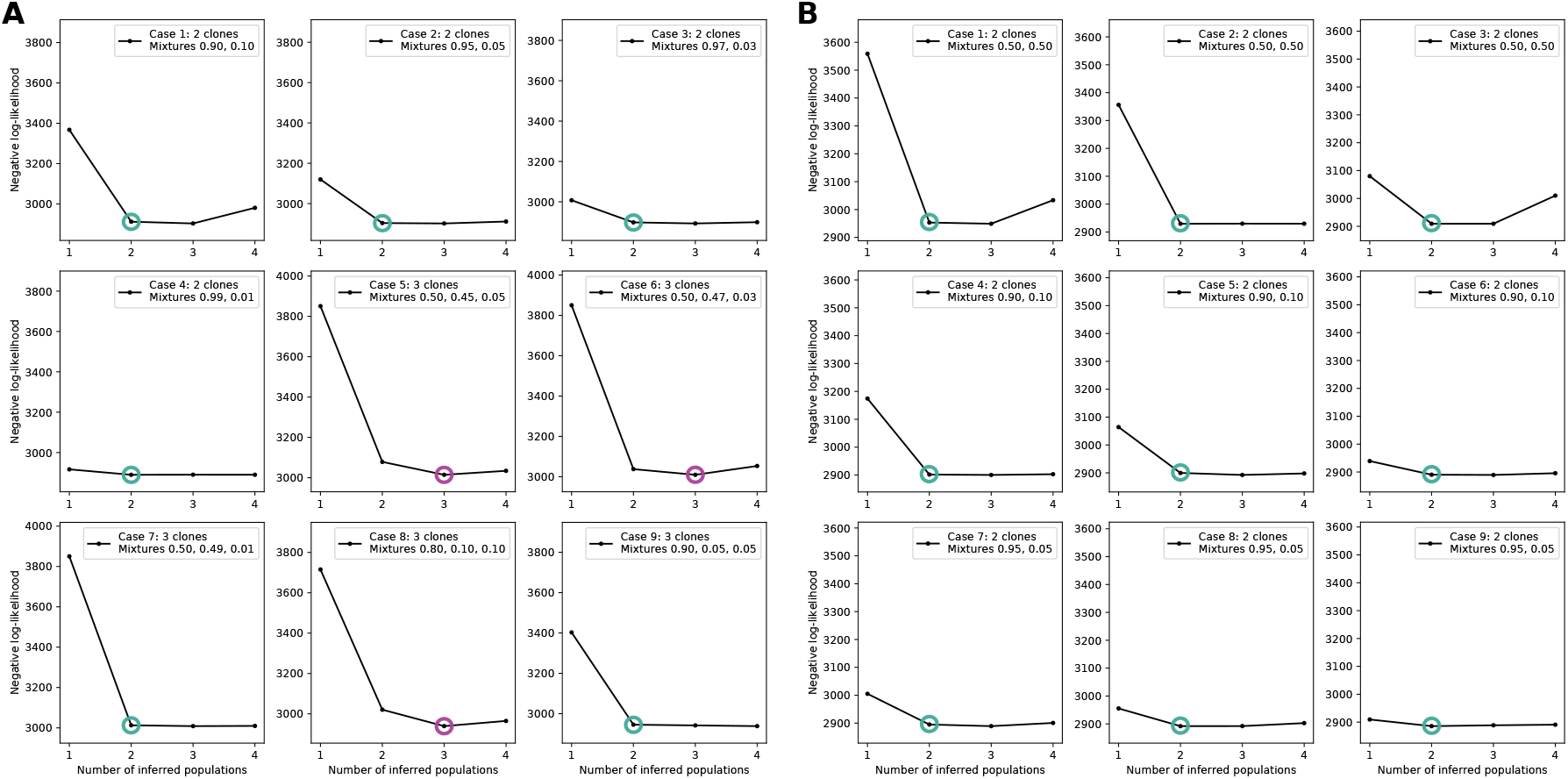
Related to Figure 7. Elbow plots showing the negative log-likelihood for all cases in Figure 7, with the selected model marked by a circle. The color of the circle also indicates the selected model: blue for 1 population, teal for 2, dark magenta for 3.

## References

1. Janiszewska, M., Stein, S., Metzger Filho, O., Eng, J., Kingston, N. L., Harper, N. W., Rye, I. H., Alečković, M., Trinh, A., Murphy, K. C. et al. (2021). The impact of tumor epithelial and microenvironmental heterogeneity on treatment responses in HER2+ breast cancer. JCI insight, 6.

2. Marusyk, A., Janiszewska, M., and Polyak, K. (2020). Intratumor heterogeneity: the rosetta stone of therapy resistance. Cancer cell, 37, 471–484.

3. Jia, D., Lu, M., Jung, K. H., Park, J. H., Yu, L., Onuchic, J. N., Kaipparettu, B. A., and Levine, H. (2019). Elucidating cancer metabolic plasticity by coupling gene regulation with metabolic pathways. Proceedings of the National Academy of Sciences, 116, 3909–3918.

4. Moldogazieva, N. T., Mokhosoev, I. M., and Terentiev, A. A. (2020). Metabolic heterogeneity of cancer cells: an interplay between HIF-1, GLUTs, and AMPK. Cancers, 12, 862.

5. Ng, C. K., Weigelt, B., A’Hern, R., Bidard, F. C., Lemetre, C., Swanton, C., Shen, R., and Reis-Filho, J. S. (2014). Predictive Performance of Microarray Gene Signatures: Impact of Tumor Heterogeneity and Multiple Mechanisms of Drug Resistance. Cancer research, 74, 2946–2961.

6. Greaves, M., and Maley, C. C. (2012). Clonal evolution in cancer. Nature, 481, 306–313.

7. Maley, C. C., Galipeau, P. C., Finley, J. C., Wongsurawat, V. J., Li, X., Sanchez, C. A., Paulson, T. G., Blount, P. L., Risques, R. A., Rabinovitch, P. S. et al. (2006). Genetic clonal diversity predicts progression to esophageal adenocarcinoma. Nature genetics, 38, 468–473.

8. Dexter, D. L., and Leith, J. T. (1986). Tumor heterogeneity and drug resistance. Journal of clinical oncology, 4, 244–257.

9. Sharma, S. V., Lee, D. Y., Li, B., Quinlan, M. P., Takahashi, F., Maheswaran, S., McDermott, U., Azizian, N., Zou, L., Fischbach, M. A. et al. (2010). A chromatin-mediated reversible drug-tolerant state in cancer cell subpopulations. Cell, 141, 69–80.

10. Knoechel, B., Roderick, J. E., Williamson, K. E., Zhu, J., Lohr, J. G., Cotton, M. J., Gillespie, S. M., Fernandez, D., Ku, M., Wang, H. et al. (2014). An epigenetic mechanism of resistance to targeted therapy in T cell acute lymphoblastic leukemia. Nature genetics, 46, 364–370.

11. Hafner, M., Niepel, M., and Sorger, P. K. (2017). Alternative drug sensitivity metrics improve preclinical cancer pharmacogenomics. Nature biotechnology, 35, 500–502.

12. Pozdeyev, N., Yoo, M., Mackie, R., Schweppe, R. E., Tan, A. C., and Haugen, B. R. (2016). Integrating heterogeneous drug sensitivity data from cancer pharmacogenomic studies. Oncotarget, 7, 51619.

13. Brooks, E. A., Galarza, S., Gencoglu, M. F., Cornelison, R. C., Munson, J. M., and Peyton, S. R. (2019). Applicability of drug response metrics for cancer studies using biomaterials. Philosophical Transactions of the Royal Society B, 374, 20180226.

14. Matulis, S. M., Gupta, V. A., Neri, P., Bahlis, N. J., Maciag, P., Leverson, J. D., Heffner, L. T., Lonial, S., Nooka, A. K., Kaufman, J. L. et al. (2019). Functional profiling of venetoclax sensitivity can predict clinical response in multiple myeloma. Leukemia, 33, 1291–1296.

15. Bonolo de Campos, C., Meurice, N., Petit, J. L., Polito, A. N., Zhu, Y. X., Wang, P., Bruins, L. A., Wang, X., Armenta, I. D. L., Darvish, S. A. et al. (2020). “Direct to Drug” screening as a precision medicine tool in multiple myeloma. Blood cancer journal, 10, 54.

16. Di Veroli, G. Y., Fornari, C., Goldlust, I., Mills, G., Koh, S. B., Bramhall, J. L., Richards, F. M., and Jodrell, D. I. (2015). An automated fitting procedure and software for dose-response curves with multiphasic features. Scientific reports, 5, 14701.

17. Muthén, B., and Shedden, K. (1999). Finite mixture modeling with mixture outcomes using the EM algorithm. Biometrics, 55, 463–469.

18. Jung, T., and Wickrama, K. A. (2008). An introduction to latent class growth analysis and growth mixture modeling. Social and personality psychology compass, 2, 302–317.

19. Hafner, M., Niepel, M., Chung, M., and Sorger, P. K. (2016). Growth rate inhibition metrics correct for confounders in measuring sensitivity to cancer drugs. Nature methods, 13, 521–527.

20. Cadena-Herrera, D., Esparza-De Lara, J. E., Ramírez-Ibañez, N. D., López-Morales, C. A., Pérez, N. O., Flores-Ortiz, L. F., and Medina-Rivero, E. (2015). Validation of three viable-cell counting methods: Manual, semi-automated, and automated. Biotechnology Reports, 7, 9–16.

21. Mumenthaler, S. M., Foo, J., Leder, K., Choi, N. C., Agus, D. B., Pao, W., Mallick, P., and Michor, F. (2011). Evolutionary modeling of combination treatment strategies to overcome resistance to tyrosine kinase inhibitors in non-small cell lung cancer. Molecular pharmaceutics, 8, 2069–2079.

22. Tadele, D. S., Robertson, J., Crispin, R., Herrera, M. C., Chlubnová, M., Piechaczyk, L., Ayuda-Durán, P., Singh, S. K., Gedde-Dahl, T., Fløisand, Y. et al. (2021). A cell competition-based small molecule screen identifies a novel compound that induces dual c-Myc depletion and p53 activation. Journal of Biological Chemistry, 296.

23. Kumar, S., Paiva, B., Anderson, K. C., Durie, B., Landgren, O., Moreau, P., Munshi, N., Lonial, S., Bladé, J., Mateos, M. V. et al. (2016). International Myeloma Working Group consensus criteria for response and minimal residual disease assessment in multiple myeloma. The lancet oncology, 17, 328–346.

24. Keats, J. J., Chesi, M., Egan, J. B., Garbitt, V. M., Palmer, S. E., Braggio, E., Van Wier, S., Blackburn, P. R., Baker, A. S., Dispenzieri, A. et al. (2012). Clonal competition with alternating dominance in multiple myeloma. Blood, The Journal of the American Society of Hematology, 120, 1067–1076.

25. Lohr, J. G., Stojanov, P., Carter, S. L., Cruz-Gordillo, P., Lawrence, M. S., Auclair, D., Sougnez, C., Knoechel, B., Gould, J., Saksena, G. et al. (2014). Widespread genetic heterogeneity in multiple myeloma: implications for targeted therapy. Cancer cell, 25, 91–101.

26. Giliberto, M., Thimiri Govinda Raj, D. B., Cremaschi, A., SkAanland, S. S., Gade, A., Tjønnfjord, G. E., Schjesvold, F., Munthe, L. A., and Taskén, K. (2022). Ex vivo drug sensitivity screening in multiple myeloma identifies drug combinations that act synergistically. Molecular Oncology, 16, 1241–1258.

27. Shi, J., Alagoz, O., Erenay, F. S., and Su, Q. (2014). A survey of optimization models on cancer chemotherapy treatment planning. Annals of Operations Research, 221, 331–356.

28. Lai, X., Geier, O. M., Fleischer, T., Garred, Ø., Borgen, E., Funke, S. W., Kumar, S., Rognes, M. E., Seierstad, T., Børresen-Dale, A. L. et al. (2019). Toward personalized computer simulation of breast cancer treatment: A multiscale pharmacokinetic and pharmacodynamic model informed by multitype patient data. Cancer research, 79, 4293–4304.

29. Swan, G. W. (1990). Role of optimal control theory in cancer chemotherapy. Mathematical biosciences, 101, 237–284.

30. He, Q., Zhu, J., Dingli, D., Foo, J., and Leder, K. Z. (2016). Optimized treatment schedules for chronic myeloid leukemia. PLoS computational biology, 12, 1005129.

31. Moulines, E., Cardoso, J. F., and Gassiat, E. Maximum likelihood for blind separation and deconvolution of noisy signals using mixture models. In 1997 IEEE international conference on acoustics, speech, and signal processing (Vol. 5, pp. 3617–3620), 1997.

32. Comon, P., and Jutten, C. (2010). Handbook of Blind Source Separation: Independent component analysis and applications (Academic press).

33. Sage, D., Prodanov, D., Tinevez, J. Y., and Schindelin, J. MIJ: making interoperability between ImageJ and Matlab possible. In ImageJ User & Developer Conference (Vol. 2426), 2012.

34. Wang, D., Fløisand, Y., Myklebust, C. V., Bürgler, S., Parente-Ribes, A., Hofgaard, P. O., Bogen, B., Tasken, K., Tjønnfjord, G. E., Schjesvold, F. et al. (2017). Autologous bone marrow Th cells can support multiple myeloma cell proliferation in vitro and in xenografted mice. Leukemia, 31, 2114–2121.

35. Riss, T. L., Moravec, R. A., Niles, A. L., Duellman, S., Benink, H. A., Worzella, T. J., and Minor, L. (2016). Cell viability assays. Assay Guidance Manual.

36. Garvey, C. M., Spiller, E., Lindsay, D., Chiang, C. T., Choi, N. C., Agus, D. B., Mallick, P., Foo, J., and Mumenthaler, S. M. (2016). A high-content image-based method for quantitatively studying context-dependent cell population dynamics. Scientific reports, 6, 1–12.

37. Celeux, G., Frühwirth-Schnatter, S., and Robert, C. P. (2019). Model selection for mixture models-perspectives and strategies. In Handbook of mixture analysis, G. Celeux, S. Frühwirth-Schnatter, and C. P. Robert, ed. (Chapman and Hall/CRC), pp. 117–154.

38. MathWorks. (2020). MATLAB Optimization Toolbox. https://se.mathworks.com/products/optimization.html.

39. Tang, M., Zhao, R., van de Velde, H., Tross, J. G., Mitsiades, C., Viselli, S., Neuwirth, R., Esseltine, D.L., Anderson, K., Ghobrial, I. M., et al. (2016). Myeloma Cell Dynamics in Response to Treatment Supports a Model of Hierarchical Differentiation and Clonal Evolution. Clinical Cancer Research, 22, 4206–4214.

40. Milo, R., Jorgensen, P., Moran, U., Weber, G., and Springer, M. (2010). BioNumbers—the database of key numbers in molecular and cell biology. Nucleic acids research, 38, D750–D753.

